# A high-resolution atlas of cattle regulatory variants and their cross-species activity in matched human cells

**DOI:** 10.64898/2026.05.06.723179

**Authors:** Rongrong Zhao, Lindsey J. Plenderleith, Tanmay Debnath, Rachel S. Owen, Ludo Pagie, Vartika Bisht, Carey Metheringham, Melissa M. Marr, Jessica Powell, Andrea Talenti, Chumeng Zhu, Edith Paxton, Kirsty Jensen, Joris van Arensbergen, Tim K. Connelley, Liam J. Morrison, Musa A. Hassan, James G. D. Prendergast

## Abstract

Identifying causal noncoding variants underlying complex traits in cattle remains challenging because high-resolution functional maps of regulatory variation are lacking. Here we combine massively parallel reporter assays with graph genomics to measure autonomous transcriptional activity from >1.5 billion DNA fragments spanning both cattle subspecies. In primary bovine cells, we assay >15 million variants and identify >150,000 expression-modulating variants enriched at cattle eQTL and GWAS loci. This enables the refinement of broad association signals to small sets of candidate functional regulatory variants. Our haplotype-aware framework captures rare, multi-allelic and tightly linked variants poorly resolved by conventional eQTL studies, and quantifies the disproportionate impact of larger variants on transcription. Furthermore, we use these data to train a deep-learning model that successfully predicts bovine promoter activity directly from sequence. Profiling the same cattle DNA in matched primary human cells reveals widespread conservation of promoter and enhancer activity, allelic effects and regulatory grammar, supporting the transfer of annotations and models across species. However, species-dependent effects are enriched in evolutionarily young sequences and p53-family motifs, highlighting the limits to simple cross-species extrapolation. Together, these data provide a high-resolution atlas of cattle regulatory variation and a framework for prioritising causal noncoding variants for cattle trait improvement.

## Introduction

Livestock sit at the intersection of global food security, climate change and important zoonotic diseases. Decades of genome-wide association studies (GWAS) in cattle have identified hundreds of loci linked to relevant traits including production, methane emissions and disease resistance^1^. However, in almost all cases the underlying causal variants remain unresolved, limiting the ability to directly target them to improve cattle via advanced breeding approaches such as genome editing or weighted polygenic trait prediction^2^. Across mammals, diverse lines of evidence suggest that noncoding regulatory variants are the primary drivers of complex traits^3–5^, meaning that pinpointing functional regulatory variants in livestock genomes is a critical step towards accelerating genetic improvement for important phenotypes.

The difficulty of identifying functional regulatory variants in the cattle genome is compounded by its extensive levels of linkage disequilibrium^6^, meaning traditional expression quantitative trait loci (eQTL) studies can generally only map larger loci and not causal variants. Moreover, the extensive complementary resources available to the human genomics community for fine-mapping, such as high-resolution maps of regulatory elements across cell types^7–9^, are at a scale that far exceed what is currently feasible to generate in livestock. However, promoters show sequence and functional conservation across mammals^10^, and although enhancer sequences evolve more rapidly, enhancer activity^11^ and regulatory logic^12^ frequently remain conserved across species. If regulatory grammar conservation across species can be determined, then the rich trove of human functional regulatory data and predictive models could potentially be leveraged to predict and prioritise causal variants in cattle and other domesticated species.

Here we use the Survey of Regulatory Elements (SuRE) massively parallel reporter assay (MPRA) approach^13^ to assay the transcriptional activity of over 1.5 billion DNA fragments from both the *Bos indicus* and *Bos taurus* cattle sub-species, spanning over 15 million genetic variants. We then test the regulatory potential of these DNA segments and variants in not only bovine but also matched human primary cell lines. By leveraging graph genomics and AI approaches, this design enables the direct identification of sequences and variants that modulate transcription in cattle, and the delineation of species-shared versus species-specific effects of the same DNA sequences. This work provides the first high-resolution catalogue of >150,000 bovine regulatory variants of this type, delivers quantitative insights into the conservation of regulatory grammar between cattle and humans, and introduces practical predictive models and data for prioritising actionable noncoding variants in livestock for trait improvement.

## Results

### A genome-wide map of autonomous regulatory activity in primary cattle cells

The SuRE MPRA approach involves fragmenting whole genomes into genomic DNA segments of a few hundred base pairs in length, then testing if these fragments can induce transcription in a promoter-less reporter plasmid^13^. In this way, it is possible to define functional transcription start sites (TSSs), spanning both promoters and enhancers, across the whole genome.

We generated SuRE genome-wide libraries from genomic DNA of two animals representing the major domesticated cattle subspecies (*Bos taurus* and *Bos indicus*), maximising the number of variant sites that can be tested downstream. Each library contained over 700 million fragments, with median lengths of 548 base pairs (bp) in *B. taurus* and 586 bp in *B. indicus*. We transfected these libraries concurrently into primary bovine aortic endothelial cells (BAOECs) in two independent biological replicates and quantified reporter initiation by sequencing the expressed barcodes, as in the original SuRE framework^14^. SuRE signal was observed to be highly reproducible between replicates (Fig. 1), enriched at transcription start sites (Fig. 1b) and correlated with promoter PRO-cap^14^ signal generated from the same cells (Pearson’s *r* > 0.7, *P* < 1 × 10^−300^; Fig. 1c), indicating that SuRE successfully captures a substantial component of endogenous cattle transcription initiation.

**Figure 1.**
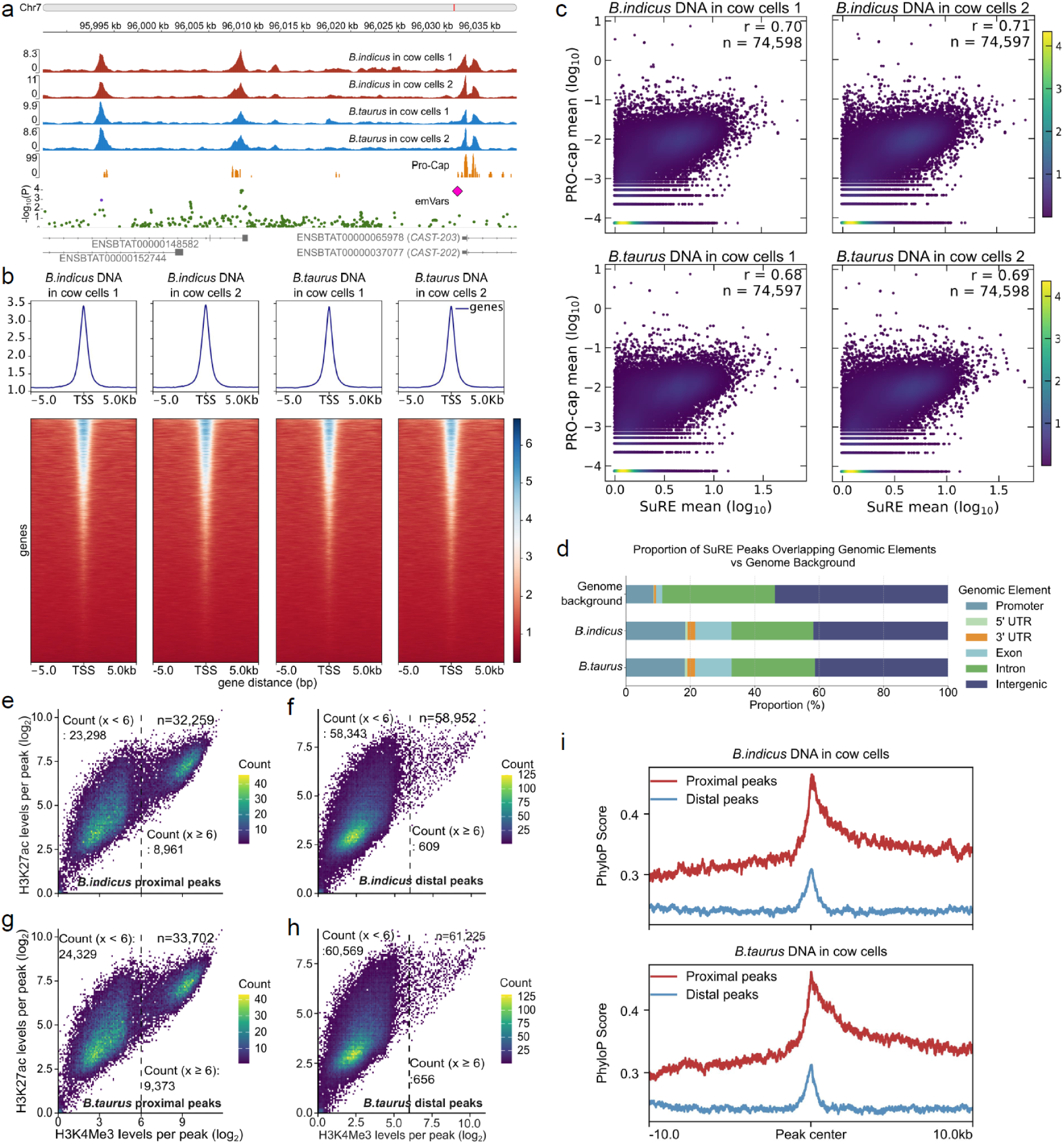
SuRE accurately captures proximal and distal regulatory elements in the cattle genome. (a) Representative SuRE and PRO-cap signal tracks at the *CAST* gene locus on chromosome 7, known to harbour an eQTL previously linked to the meat tenderness trait^15^. The profiles demonstrate that SuRE accurately identifies both proximal and distal regulatory elements with high concordance with endogenous PRO-cap transcription signals. The track of expression-modulating variants (emVars) discussed below is shown in green. The lead emVar at the CAST TSS is indicated by an enlarged purple diamond. (b) Average SuRE signal profiles (top) and heatmaps (bottom) illustrating signal enrichment near cattle transcription start sites (TSSs). The x-axis spans 10 kb regions centred on TSSs; heatmaps represent individual genes ranked by SuRE signal intensity. (c) Correlation between SuRE mean activity and PRO-cap mean signal at cattle TSSs for *B. indicus* (*r* = 0.70-0.71) and *B. taurus* (*r* = 0.68-0.69) DNA across two independent biological replicates. (d) Proportion of SuRE-called peaks overlapping known cattle genomic elements compared to the genome-wide background distribution. Where relevant, peaks were assigned to one group only (see methods). (e-h) Density scatter plots comparing H3K4me3 and H3K27ac levels per SuRE peak for *B. indicus* (e, f) and *B. taurus* (g, h) DNA. Peaks are subdivided into proximal elements (e, g), which show high enrichment for both histone marks, and distal elements (f, h), which are primarily enriched for the enhancer-associated H3K27ac mark only. (i) Evolutionary conservation profiles (PhyloP scores) centred on SuRE proximal and distal peaks for *B. indicus* and *B. taurus*, highlighting elevated conservation at the centres of both distal and proximal peaks.

Peak calling identified 94,927 candidate regulatory elements in the *B. taurus* library and 91,211 in the *B. indicus* library. Approximately 18% overlapped annotated promoters, representing a clear enrichment relative to the genome-wide background (8.6%; Χ^2^ = 3784.5, *P* < 1 × 10^−300^; Fig. 1d). Proximal peaks (≤10 kb to nearest TSS), were enriched for both H3K27ac and H3K4me3, whereas distal peaks (>10 kb to nearest TSS) showed high H3K27ac with relatively low H3K4me3 (Fig. 1e–h), consistent with promoter-like and enhancer-like chromatin states, respectively. Both proximal and distal peaks were enriched at regions of elevated sequence conservation across species (Fig. 1i), also consistent with conserved functional elements. Together, these results establish that this SuRE dataset provides a reproducible, high-resolution, genome-wide map of autonomous regulatory activity in primary cattle cells.

### Graph-based analysis increases regulatory variant recovery and power

As SuRE assays many overlapping genomic fragments, it can be used to fine-map regulatory variants by testing whether fragments that carry different alleles drive different levels of transcription. Doing this at scale requires two considerations: accurate mapping of SuRE fragments back to the genome, and reliable classification of fragments by the allele they carry at each variant. In cattle, the high variant density and divergence between *B. taurus* and *B. indicus* can introduce reference bias when aligning fragment reads to a single linear (*B. taurus*) reference^16^, reducing mapping rates and allele coverage, a particular issue for the indicine-derived library and for longer variants.

We therefore analysed the SuRE libraries using a novel graph genome workflow (Fig. 2a), constructed from haplotype-resolved assemblies of the two DNA donor cattle together with the *B. taurus* reference genome^17^. With the aim of reducing reference mapping bias and, importantly, enabling inference of internal alleles on DNA fragment sections not directly covered by sequencing reads, increasing effective allele coverage for downstream analyses (Fig. 2a).

**Figure 2.**
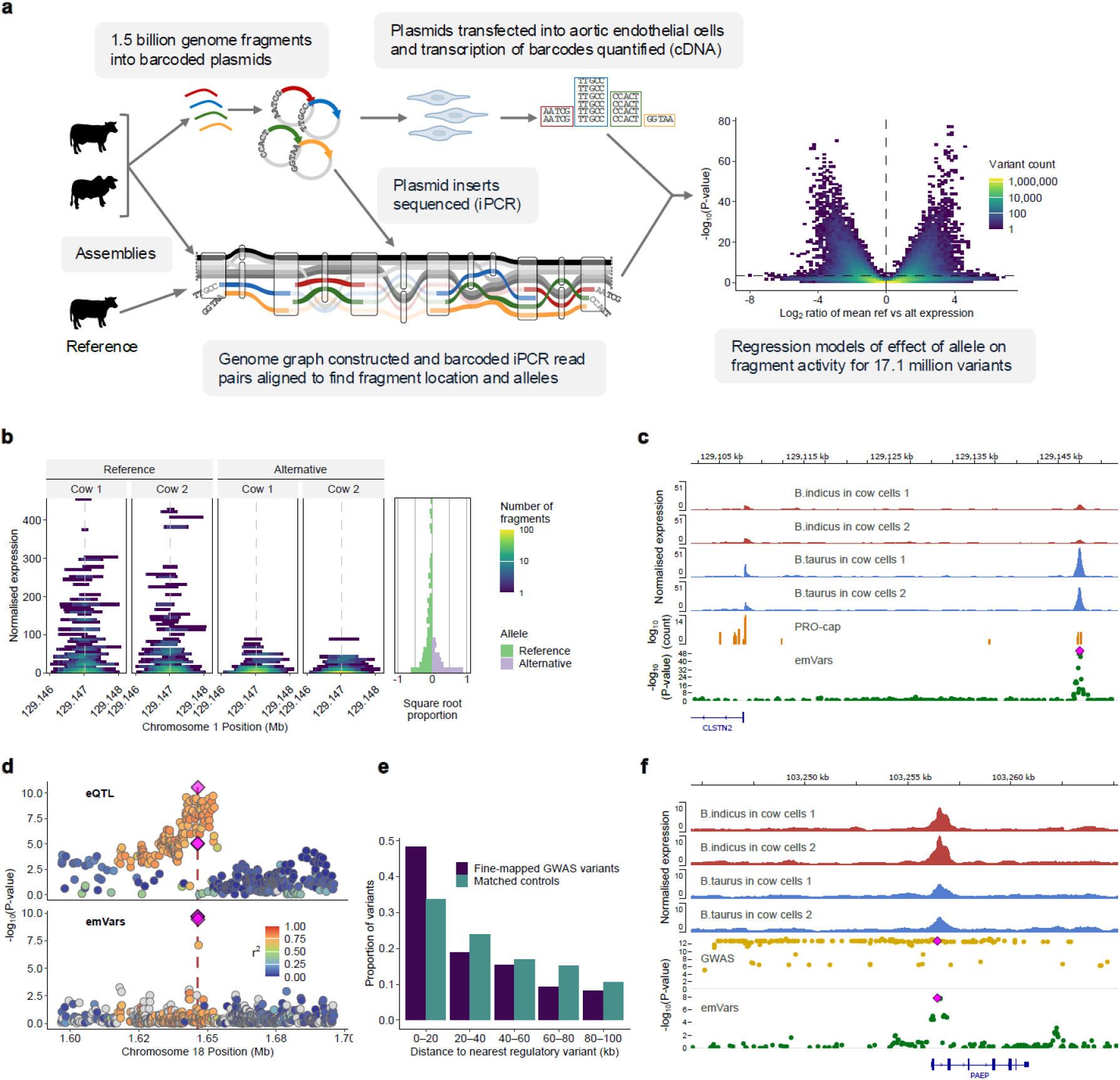
Graph-based SuRE profiling expands regulatory variant detection and refines trait loci. (a) Schematic of the haplotype-aware genome graph processing workflow (left) and the resulting identification of expression-modulating variants. The square-binned density plot (right) displays all tested biallelic variants. The x-axis shows the log2-transformed expression ratio between reference and alternative alleles, and the y-axis shows the -log10-transformed P-value from the allele-specific expression model. Colour intensity reflects the variant count per bin. The horizontal dashed line denotes the FDR ≤ 0.05 significance threshold. (b) Allele-specific transcriptional activity at a distal regulatory variant (1:129147515:G:A) upstream of *CLSTN2* linked to carcass traits. The main panels plot individual DNA fragments according to their genomic location (x-axis), normalized SuRE score (y-axis), and the allele they carry (reference or alternative) across the two biological replicates. Colour intensity indicates overlapping fragment density. The mirrored bar chart (right) summarises the allele-specific expression distribution, showing the square root of the fragment proportion in the given expression bin for the reference (green) and alternative (purple) alleles. (c) Genomic tracks at the *CLSTN2* locus showing SuRE signal (red/blue), endogenous PRO-cap transcription initiation (orange), and emVar significance (green points). The focal 1:129147515:G:A variant is highlighted in magenta. (d) Refinement of broad eQTL associations using SuRE data. The upper plot displays cattle GTEx eQTL associations (-log10-transformed nominal P-values) for the *FCSK* gene in adipose tissue, which include 99 variants passing the threshold for *cis*-eQTL significance; the lower plot shows the corresponding SuRE emVar significance for variants in the same region, of which four pass the threshold for significance (highlighted by magenta diamonds). Variants are coloured by their pairwise linkage disequilibrium (r^2^) relative to the focal SuRE variant (vertical dashed line). (e) Distance distribution of fine-mapped GWAS causal variants (dark blue) and matched control variants (teal) relative to the nearest SuRE regulatory element. Control variants were matched for chromosome and *B. taurus* minor allele frequency. Fine-mapped variants localise significantly closer to emVars than the matched controls (Mann-Whitney U test, P = 9.6x10^-10^). (f) SuRE pinpoints candidate causal variants at the *PAEP* gene locus. Tracks show normalized SuRE signal (red and blue), SuRE emVars (green points) and GWAS variants associated with milk saturated fatty acid levels^20^ (brown points). The most significant emVar is highlighted in magenta across both variant tracks.

Using this graph-aware pipeline, we identified and screened 14.8 million biallelic single nucleotide variants (SNVs) segregating within or between the two donor genomes for allele-specific SuRE activity. At an FDR threshold of 0.05, we identified 140,794 expression-modulating variants (emVars). In contrast, using an analogous linear-reference workflow adapted from prior SuRE analyses^13^ detected 76,900 emVars among 12.1 million tested SNVs, indicating that this graph-based alignment and phased allele inference increased both the number of testable variants and power to detect allele-specific effects at a fixed FDR.

Among the putative regulatory variants resolved by this approach is the 1:129147515:G:A variant (Fig. 2b), located at a putative distal element upstream of the *CLSTN2* gene. This locus has previously been associated with carcass traits in *B. indicus* cattle^18^ and the regulatory activity of the site is corroborated by endogenous PRO-cap data (Fig. 2c).

### emVars refine endogenous eQTL and GWAS signals

Across cattle GTEx^19^ tissues, emVars consistently showed stronger eQTL signal than assayed variants not called as emVars (one-sided Wilcoxon rank-sum test, *P* < 7 × 10^−4^ in each tissue; Supplementary Fig. 1, Supplementary Table 1). The higher resolution of SuRE was observed to successfully refine broad eQTL association intervals to smaller sets of candidate regulatory variants at individual loci (Fig. 2d). Notably many of the identified emVars would be excluded by allele frequency filters commonly used in eQTL studies; with 37,119 and 56,827 significant emVars having *B. taurus* allele frequencies below commonly used thresholds of 1% and 5%, respectively. Thus, up to 40% of emVars would be undetectable in conventional eQTL analyses.

The observation that emVars preferentially coincide with eQTL signals across many tissues suggests that a substantial fraction of SuRE-detected regulatory effects are relevant beyond the assayed endothelial cell context. This included their enrichment at mammary eQTLs (one-sided Wilcoxon rank-sum P = 1.52x10^-17^, Supplementary Table 1), consistent with potential relevance to fine-mapping dairy phenotypes. Supporting this, emVars were enriched near fine-mapped GWAS variants for dairy traits (Fig. 2e). For example, at the *PAEP* locus, which encodes the primary whey protein in milk that has been linked to multiple dairy traits, including saturated fatty acid levels^20^, SuRE refines a broader GWAS interval to a small set of promoter emVars (Fig. 2f), highlighting concrete candidate causal variants for downstream follow-up.

### Larger and multi-allelic variants add substantial regulatory signal

Most functional maps of regulatory variation in cattle, including eQTL studies, are biased towards SNVs due to the difficulty of genotyping larger variants. However, cattle genomes harbour abundant larger complex variants, and there is substantial interest in the idea that these variants may disproportionately underpin important cattle traits^21^. SuRE combined with graph-based genotyping provides an opportunity to test this directly at scale.

Beyond SNVs, the genome graph contained ∼2 million other biallelic variants: predominantly indels (n = 1,523,290), with the remainder comprising substitutions of equal (n = 388,898) or unequal (n = 43,094) length. 16,111 variants affected ≥50 nucleotides (15,779 indels and 332 substitutions), referred to here as SV-class variants. Applying analogous allele-specific models to this non-SNV set identified 18,934 functional non-SNV regulatory variants, including 422 SV-class variants (FDR ≤ 0.05; Fig. 3a). The differential transcription induced by fragments overlapping the 11:30920596 16 bp insertion in the third exon of *GTF2A1L* is shown in Fig. 3b (Supplementary Fig. 2). The probability of regulatory activity was observed to increase strongly with variant size (Fig. 3c). Variants >100 bp were more than four-fold as likely to be regulatory than 1-nt indels (Chi-squared test P < 2.2x10^-16^), and the same trend was observed for allele length change (Fig. 3d, Chi-squared test P < 2.2x10^-16^ for 1-nt vs 100-nt for both insertions and deletions).

**Figure 3.**
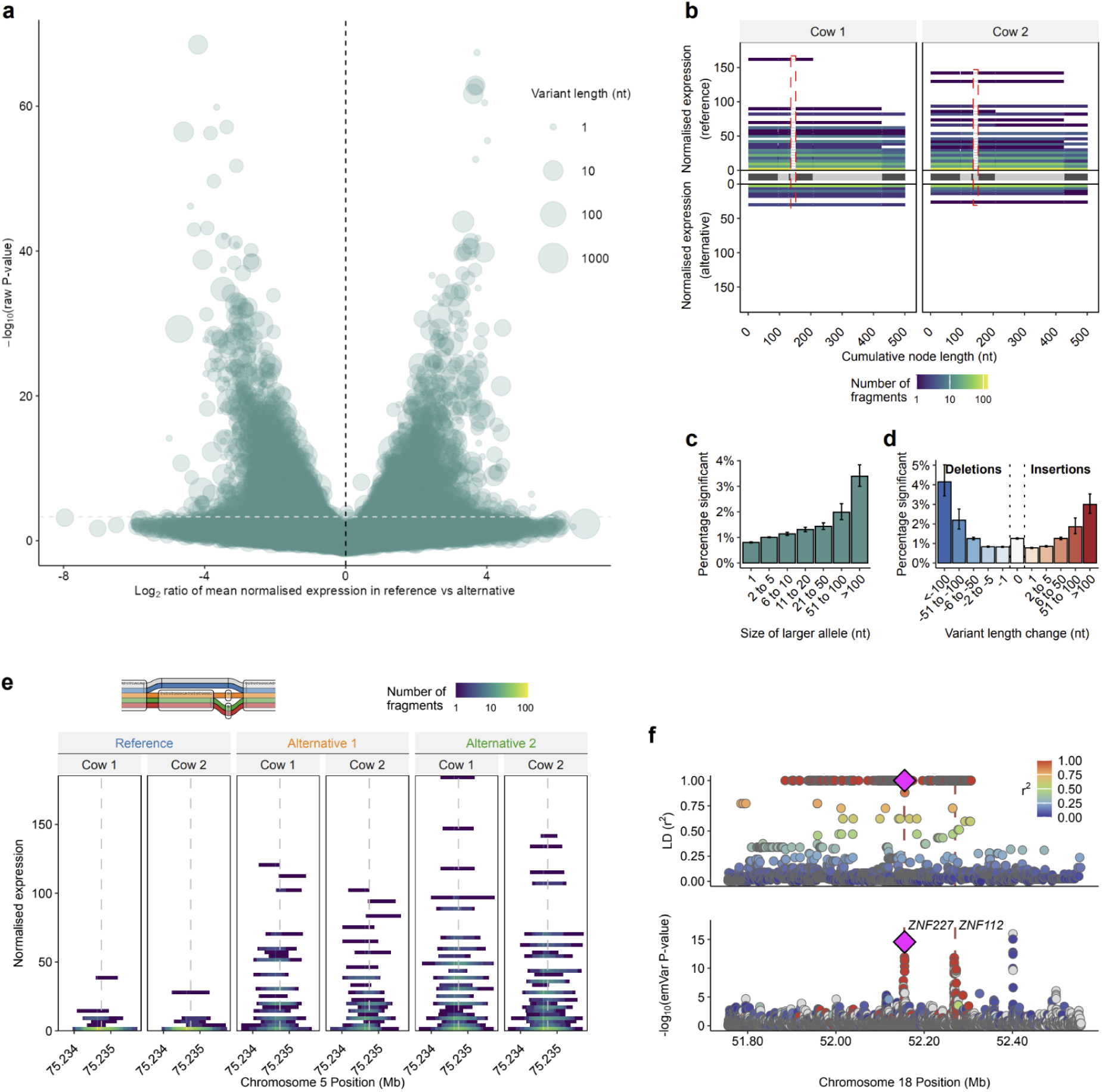
SuRE allows the regulatory potential of more complex genetic variation to be explored. (a) Volcano plot of biallelic non-SNV emVars. The x-axis shows the log2-transformed ratio of the normalised SuRE score between fragments carrying the reference and alternative alleles, and the y-axis shows the -log10-transformed P-value. Bubble size denotes the length of the larger allele in nucleotides (nt). The horizontal dashed line indicates an FDR threshold of ≤ 0.05. (b) Allele-specific transcriptional activity of fragments overlapping a 16-nt insertion on chromosome 11 (11:30920596; FDR = 1.78x10^-16^). Fragments carrying the reference (top) and alternative (bottom) alleles are plotted by replicate across the genome graph. Colour intensity reflects fragment density per expression bin. The x-axis shows progression through the graph, with nodes in the graph shown as rectangles of alternating shades of grey. Nodes which are skipped by all fragments are shown by a thin grey line. The dashed red rectangle highlights the graph node representing the insertion, which is visited exclusively by alternative allele fragments. P-values for adjacent emVars are shown in Supplementary Fig. 2. (c-d) The probability of a non-SNV exhibiting regulatory activity scales with variant size. Bar charts display the percentage of tested variants that were significant binned by (c) the absolute size of the larger allele, and (d) the net length change between alleles (deletions in blue, insertions in red). Error bars represent 95% confidence intervals. The “0” bin in (d) denotes complex substitutions where allele lengths are equal (range: 2–380 nt). (d) Allele-specific expression across biological replicates for a multi-allelic variant on chromosome 5 (5:75234745; FDR = 4.75x10^-41^), located 21 kb upstream of the lncRNA gene ENSBTAG00000062845. Normalised expression is shown for each allele across replicates. The genome graph structure at this location is shown above, with colours matching the allele labels. Where two haplotypes had the same path, the colour of one representative was randomly chosen. (f) SuRE data allows resolution of multiple independent regulatory variants within regions with perfect LD. Resolution of independent regulatory signals within a region of high linkage disequilibrium (LD) near the ZNF227 and ZNF112 transcription start sites (vertical red lines). The top panel shows pairwise LD (r^2^) in a Holstein-Friesian cohort relative to the lead emVar (magenta diamond). The bottom panel displays the corresponding SuRE regulatory significance (-log10P-value). Variants are coloured by their LD to the lead variant; grey denotes tested variants in the SuRE analysis absent from the Holstein-Friesian dataset.

Beyond biallelic sites, the graph resolved 304,394 multi-allelic variants (including 23,526 SNVs with more than one alternative allele) segregating between the donor genomes. Testing these revealed 3,745 significant regulatory variants (323 SNVs and 3,422 complex variants; FDR ≤ 0.05, Fig. 3e, Supplementary Fig. 3), including a significant indel at the *CAST* locus shown in Fig. 1a (FDR = 0.018), previously linked to meat tenderness^15^. Collectively, these findings highlight a substantial pool of functional regulatory variation driven by complex and multi-allelic variants that are frequently overlooked in standard studies.

### Distinct regulatory effects within haplotypes

Effective genomic selection for trait improvement in livestock depends on the ability to select for different combinations of functional, trait-associated variants. When functional variants are in perfect linkage disequilibrium (LD) in a population and/or in close physical proximity, selecting for them independently is not possible, which can constrain trait improvement and adversely impact other phenotypes. SuRE data allows us to evaluate the potential scale of this issue in cattle. To assess the potential impact of LD between functional variants on trait improvement, we first identified pairs of emVars that were each significantly associated with regulatory activity in SuRE and were also in perfect LD with each other in a cohort of Holstein–Friesian cattle. We restricted this analysis to emVar pairs separated by at least 5 kb ensuring that the two variants were captured on distinct SuRE fragments and therefore assayed independently. We identified 10,937 distinct emVars among these 28,103 perfectly linked emVar pairs (Fig. 3f, Supplementary Fig. 4). This indicates that even when conservatively restricting to variants >5kb apart and in perfect LD, a substantial fraction of functional regulatory variants (7.8% of all emVars) can not be readily selected for independently. Selecting for one will necessarily alter the frequency of another.

We next examined significant emVars within 200 bp of one another that were therefore frequently observed on the same fragments. Although for most pairs independent regulatory effects could not be resolved because of complete linkage or nearby untestable variants (see methods, n = 86,879), 5,349 pairs remained that occurred in multiple haplotype combinations. Conditional modelling of fragments spanning both variants identified 388 lead variants that occurred in pairs where both sites showed evidence of contributing independent effects on regulatory activity. The proportion of variants with separable effects was observed to increase with the distance between them (one-sided Wilcoxon test p=0.014, Supplementary Fig. 5).

Together, these analyses highlight how thousands of variants with independent regulatory effects are tightly linked in the cattle genome, with important implications for the limits of genomic selection.

### Regulatory variants are linked to subspecies differentiation

A limitation of association-based eQTL mapping is that the ability to detect regulatory effects is tightly coupled to the variant’s allele frequency in the sampled population. In SuRE, detection instead depends primarily on whether both alleles are present in the donor genomes and on fragment coverage, allowing direct interrogation of how allele frequency and population differentiation relate to regulatory function.

We modelled the probability that a tested SNV is an emVar as a function of derived allele frequency (DAF) within each subspecies, controlling for local transcription initiation levels (PRO-cap), sampled animal genotypes and local variant density. Higher DAF in either sub-species was associated with greater odds that a tested SNV is regulatory. For example, an SNV with a DAF = 1 in *B. indicus* had 37% higher odds of being an emVar than an SNV with a DAF = 0 (OR = 1.37, 95% CI 1.33–1.41; Supplementary Table 2). A significant negative interaction between indicine and taurine DAF indicated that regulatory variants are most enriched among sites with strongly differentiated allele frequencies between the subspecies (Supplementary Fig. 6). This includes the distal variant upstream of the *CLSTN2* gene (Fig. 2c) that has an alternative allele frequency of 0.67 in *B. indicus* but 0.03 in *B. taurus*. These data therefore provide not only a functional resource for variant prioritisation, but also a framework for studying how selection and domestication have shaped regulatory architecture in cattle.

### A SuRE-trained sequence model predicts promoter activity

We next explored the ability to predict cattle promoter element activity directly from DNA sequence. To do this, we trained a convolutional neural network, EvaReg, on promoter-level SuRE scores obtained by aggregating fragment activity across fixed 600 bp windows around 35,351 annotated TSSs in *B. taurus* (Fig. 4a). On held-out chromosomes, predicted and observed promoter SuRE scores were strongly correlated (Pearson’s *r* = 0.77, Fig. 4b), and EvaReg predictions also correlated with endogenous promoter initiation activity measured by PRO-cap (Spearman’s *r* = 0.68; Supplementary Fig. 7). The model also recovered known activating motifs (Fig. 4c) and recapitulated the position-dependent effects of transcription factor motifs observed in humans^22^ (Fig. 4d), supporting the idea the model had successfully characterised underlying regulatory grammar.

**Figure 4.**
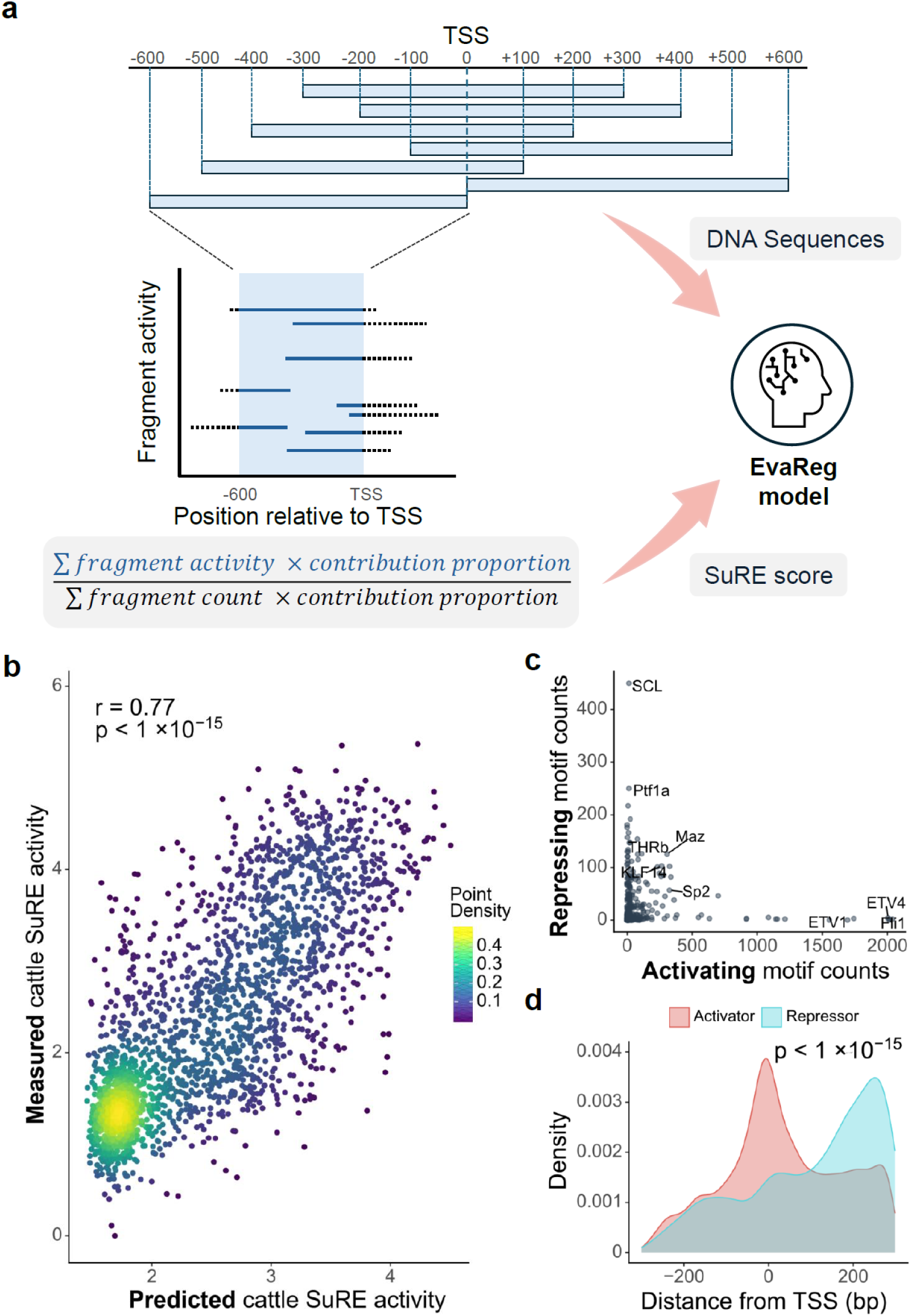
EvaReg predicts regulatory activity and learns regulatory grammar. (a) Schematic overview of the EvaReg training framework. The model learns to predict regulatory activity by evaluating 600 bp sequence windows around transcription start sites (TSSs). The SuRE score used for model training is calculated by summing the weighted contributions of each fragment that overlaps each of these windows. (b) Correlation between measured and predicted SuRE activity for promoter windows on held-out chromosomes (chromosomes 8 and 13) in *B. taurus* (Pearson’s *r*=0.77; p<1×10^-15^). (c) Scatter plot showing the total counts of repressive and activating motif instances. (d) Positional distribution of motifs identified by EvaReg as having an activating (red) or repressing (blue) function, plotted by distance from the respective TSS (Kolmogorov-Smirnov test; p<1×10^-15^).

Although EvaReg is trained on baseline promoter activity, and not fine-tuned for variant effect prediction, for SNVs within promoter windows, the absolute change in predicted score between reference and alternative alleles was larger for SuRE-significant promoter variants than for variants with no evidence of regulatory effect, and the direction of predicted change more often matched the observed direction of SuRE allelic effect than expected by chance (Supplementary Fig. 8). EvaReg consequently provides a potential extensible framework for sequence-based prioritisation of regulatory variants in cattle.

### Widespread conservation of cattle regulatory element and variant activity in human cells

An open question is the extent to which DNA-encoded regulatory control is conserved across cattle and humans. If DNA sequences largely have the same regulatory effect across matched cells from different species, this would open up the potential of leveraging the substantial human resources, such as sequence-to-function AI models, to inform cattle research. To test this directly, we transfected the same cattle SuRE libraries into primary human aortic endothelial cells in two independent experiments and quantified autonomous transcription initiation.

At the level of regulatory elements, both promoter-proximal and -distal cattle regulatory sequences identified in the cattle cells were observed to largely retain the capacity to autonomously initiate transcription in human cells (Fig. 5a, b). At the level of genetic variants, we repeated allele-specific SuRE analyses in the cattle-DNA-in-human dataset to identify 8,123 emVars in human cells. Of these, 23% (1,885) had also been identified as emVars in cattle cells (FDR ≤ 0.05; Methods). An example is the distal variant upstream of the *CLSTN2* gene (Fig. 2b, c) that showed consistent regulatory effects in both species (Fig. 5c). Among shared emVars, discordance in the direction of effect was rare; only six variants exhibited opposite directions of effect in cattle versus human cells. More broadly, across variants significant in cattle cells, allelic effects were predominantly directionally concordant even if not significant in human cells, with approximately 2.7-fold more sites showing the same direction of effect in cattle and human cells than showing opposing directions (Fig. 5d).

**Figure 5.**
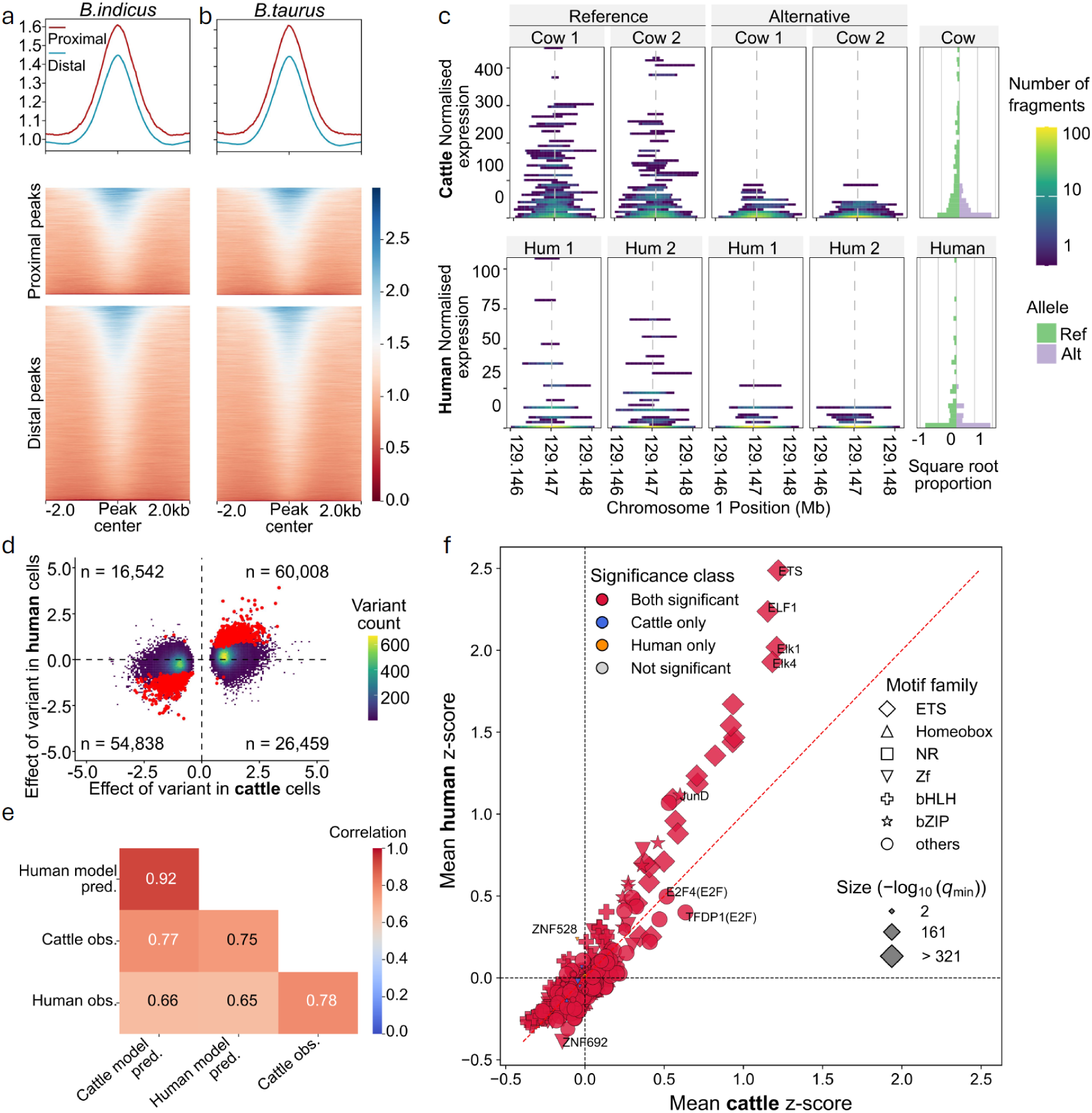
Conservation of cattle regulatory activity and grammar in human cells. (a, b) SuRE activity measured in human cells at the regulatory elements identified in cattle cells among (a) *B. indicus* and (b) *B. taurus* DNA. Average signal centred on proximal and distal peaks shows that both promoter-proximal and -distal cattle regulatory elements retain the ability to autonomously initiate transcription in human cells. Heatmaps below show signal across individual peaks, ordered by activity. (c) Conserved allele-specific regulatory activity at the distal variant upstream of *CLSTN2*. Normalized expression for the reference (Ref) and alternative (Alt) alleles is shown across replicates in cattle (top) and human cells (bottom), alongside the square root proportion of fragments in each activity bin (right). (d) Cross-species comparison of variant effect sizes for significant cattle emVars (FDR ≤ 0.05). Quadrants separate concordant (top-right, bottom-left) and discordant (top-left, bottom-right) allelic effects, with the number of variants in each quadrant indicated. Among the variants significant in both species (red points), counts per quadrant are: top-right (n = 927), bottom-left (n = 952), top-left (n = 3), and bottom-right (n = 3). (e) Correlation of predicted and observed promoter activity across species. EvaReg models independently trained on cattle and human SuRE data were evaluated on held-out cattle promoter sequences and compared to their corresponding predicted (pred.) or measured (obs.) activity. (f) Comparison of transcription factor motif importance scores inferred from EvaReg models trained in cattle versus human cells. The strong cross-species correlation indicates that the models learn broadly similar regulatory grammar. The importance scores measure the relative importance of each base as compared to the other alternative bases to the prediction. For a given motif occurrence, the z-score normalised importance scores were averaged across its bases. Point size reflects the significance of the motif difference from the distribution of background promoter bases. qmin corresponds to the lower of the q values across the human and cattle cell results. For motifs with qmin values of 0, their − log_10_(*q*) was set to 321.

This conservation of regulatory effects extended to sequence-based models. EvaReg models trained independently on cattle-cell and human-cell SuRE promoter data produced highly correlated predictions for held-out cattle promoter sequences, and showed comparable correspondence to measured SuRE activity in each species (Fig. 5e). Motif importance and position-dependent effects inferred from the two models were likewise broadly consistent (Fig. 5f and Supplementary Fig. 9), indicating that both models learn similar regulatory features despite being trained on SuRE data obtained in cells from different species. Taken together, these results indicate that a substantial component of regulatory grammar is conserved between human and cattle.

### Regulatory elements active specifically in human cells localise to cattle-specific sequences

To characterise any exceptions to the broadly conserved activity described above, we called SuRE peaks in the cattle-DNA-in-human cells dataset using the same procedure as for bovine cells, identifying 19,716 peaks for *B. taurus* DNA and 19,116 peaks for *B. indicus* DNA. We then stratified peaks in promoter-proximal and distal classes by distance to the nearest TSS as before, and classified each peak as cattle-specific, human-specific or shared depending on whether it was detected in bovine cells, human cells or both (see Methods, Supplementary Fig. 10).

Human-cell-specific proximal peaks were centred on strongly conserved sequence (Fig. 6a), consistent with evolutionary constraint at promoter-associated elements. In contrast, human-cell-specific distal peaks exhibited a relative reduction in sequence conservation at the peak centre relative to flanking sequence (Fig. 6b), a pattern that differed from the distal peaks identified in bovine cells. Human-cell-specific distal peaks were observed to more likely overlap LINE/L1 elements than distal peaks detected specifically in bovine cells (FDR < 0.05; Supplementary Fig. 11), consistent with evidence that LINE/L1-derived sequences contribute to evolutionarily dynamic regulatory activity in mammals^23^. However, the relative reduction in conservation persisted after excluding peaks overlapping any annotated repeat (Supplementary Fig. 12), indicating that repeat overlap alone does not fully explain this divergent conservation profile. Human-cell-specific peaks, and in particular distal peaks, were substantially less likely to have a detectable homologous region in the human genome. Only 33% of human-cell-specific distal peaks had an identifiable orthologous region in the human genome, as opposed to 52% of cattle-cell-specific distal peaks (two-tailed Chi-squared test P < 2.2x10^-16^, Supplementary Table 3).

**Figure 6.**
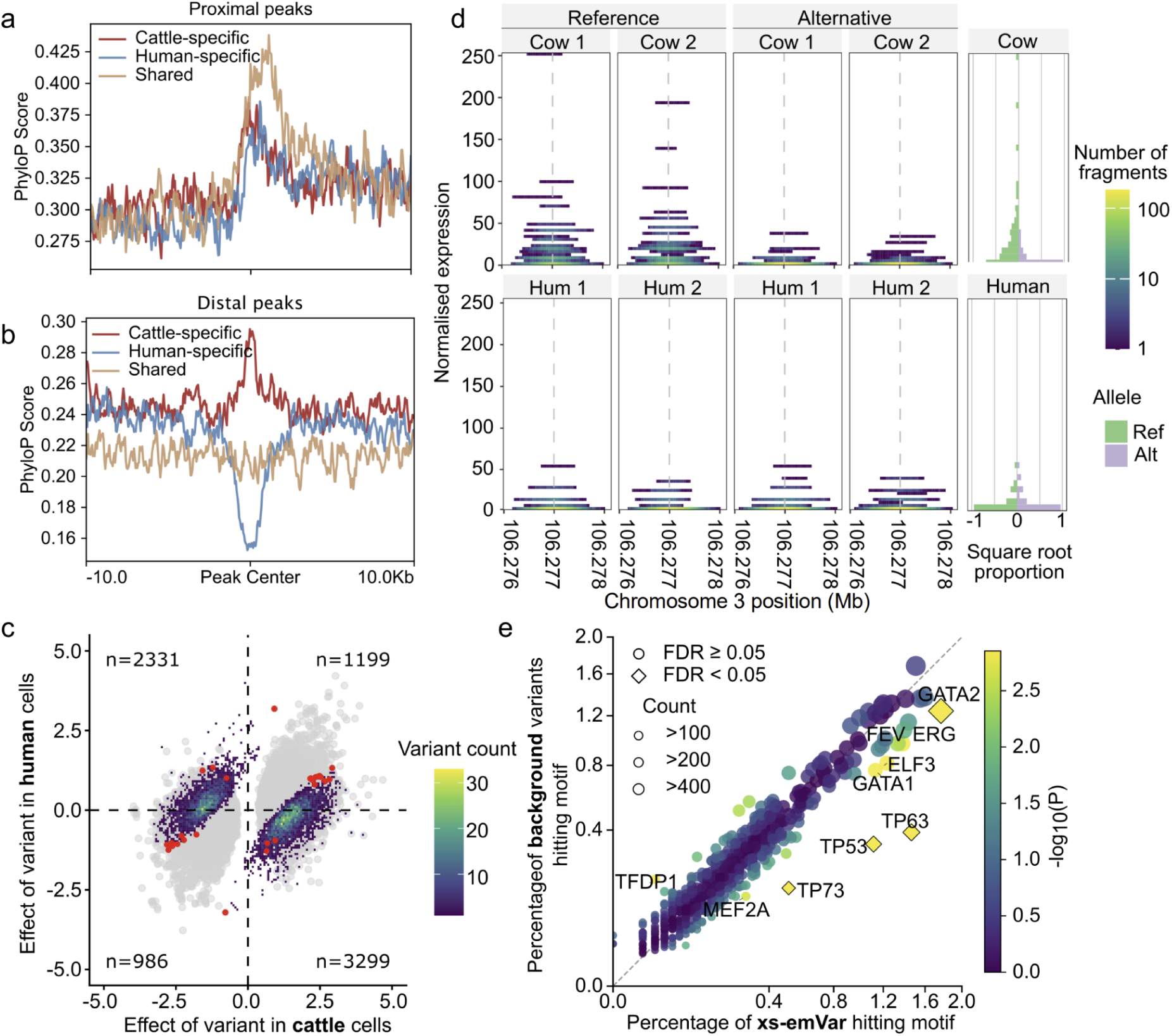
Species-dependent regulatory elements and variants exhibit distinct evolutionary and motif signatures. (a, b) Evolutionary conservation profiles (PhyloP scores) centered on SuRE proximal (a) and distal (b) peaks, stratified into cattle-specific, human-specific, and shared regulatory elements. While proximal elements broadly exhibit elevated conservation at the peak centre, distal peaks show more variable conservation patterns, including reduced conservation at the peak center for human-specific peaks relative to flanking regions. (c) Comparison of variant effects in cattle and human cells for variants with significantly different allelic effects between species (xs-emVars; FDR ≤ 0.05). xs-emVars are coloured by binned variant density and overlaid onto background variants with significant regulatory effects in cattle (grey points). Variants significant in both cattle and human cells (FDR ≤ 0.05) are highlighted in red; within this subset, quadrant counts are: top-right (n = 13), bottom-left (n = 9), top-left (n = 3), and bottom-right (n = 3). Quadrant counts for all xs-emVars are shown in the plot. (d) Representative xs-emVar on chromosome 3 demonstrating strong species-dependent regulatory activity. Normalized expression for the reference (Ref) and alternative (Alt) alleles is shown across biological replicates in cattle (top) and human (bottom) cells, alongside aggregate fragment proportion distributions (right). (e) Enrichment of transcription factor motif disruption among xs-emVars relative to background cattle emVars. Each point represents a motif, plotted by the proportion of xs-emVars versus background variants predicted to disrupt them. Colour indicates statistical significance (-log10P; two-sided Fisher’s exact test) and point size reflects variant count; significance was assessed using Benjamini-Hochberg FDR correction. Motifs from the TP53 family (TP53, TP63, TP73) and GATA factors (including GATA2) are significantly enriched among xs-emVars (FDR < 0.05).

Likewise, the 6,238 variants exhibiting regulatory effects only in human cells were less likely to fall within cattle genomic segments with a readily identifiable orthologous alignment to the human reference genome than emVars detected only in cattle cells (30% vs 43%, respectively, two-tailed Chi-squared test P < 2.2x10^-16^). A similar pattern was observed when assessing orthology to pig (31% vs 44%, P < 2.2x10^-16^). Together, these results suggest that a substantial subset of regulatory activity detected only in human cells arises from evolutionarily young or lineage-specific cattle sequence that lacks clear orthology in other mammals yet still contains binding sites capable of driving transcription in a human cellular environment.

### Cross-species QTLs are enriched for disruption of p53-family transcription factor motifs

To more explicitly identify variants with species-dependent effects, we extended the emVar model to test for species x allele interactions. In total 7,815 variants showed evidence of significantly different allelic effects between cattle and human cells (Fig. 6c), including both direction reversals (5,630 variants) and shared-direction effects with differences in magnitude (2,185 variants). The majority of these showed regulatory effects in cattle but not human cells. We refer to these as cross-species emVars (xs-emVar), an example of which is shown in Fig. 6d. To investigate potential mechanisms underlying these differences, we compared motif disruption patterns between xs-emVars and cattle emVars that were not also classified as xs-emVars (see Methods). xs-emVars were significantly enriched for variants predicted to disrupt TP53-family transcription factor motifs, including TP53, TP63 and TP73, as well as the GATA2 motif (Fig. 6e). Most other motifs showed similar disruption frequencies between xs-emVars and cattle emVars, suggesting that species-specific regulatory divergence may be partly driven by differences at certain motif classes with implications for lifting regulatory features and models between species at these locations.

## Discussion

The widespread use of genomic estimated breeding values (gEBV)^24,25^ and the emerging application of genome editing^26^ provide direct routes for translating fine-mapped regulatory variation into practical trait improvement in livestock. However, the identification of specific causal variants at most cattle trait loci has historically been hindered by long-range linkage disequilibrium and a lack of comprehensive functional maps. To overcome these limitations, we generated a high-resolution catalogue of cattle regulatory elements and fine-mapped variants that modulate transcription in unprecedented detail. Beyond the prevailing focus on short biallelic variants, our results highlight and quantify the functional relevance of understudied classes of variation, including long alleles and multi-allelic sites.

It is important to note that while we demonstrate the ability to resolve compact regulatory haplotypes harbouring multiple independent effects, when nearby variants are in perfect LD this effectively bounds the mapping resolution to a few hundred bases. In such cases, further targeted MPRA or genome editing would be required to disentangle the precise causal variant. However, to achieve meaningful improvements in gEBV accuracy and portability^27,28^, resolving regulatory variants down to this level of resolution may often be unnecessary, as recombination events within such small intervals will be rare.

Previous large-scale MPRA studies have predominantly relied on immortalised human cancer cell lines^13,29,30^, and predominantly a single biological replicate per tumour type. However, these cell lines are known to exhibit expression profiles that are distinct from primary tissues^31^, and the traits they model hold little relevance for livestock genetics. Consequently, a defining advantage of our study is the use of multiple, matching primary cell lines, which provide a more physiologically relevant context and establish a more reliable baseline for cross-species comparisons.

As SuRE measures autonomous transcription initiation from episomal DNA, it does share key limitations with other MPRA approaches; including that it does not capture the effects of native chromatin environment and long-range enhancer–promoter interactions. Consequently, SuRE reports the intrinsic regulatory potential of a DNA segment, and allele-specific effects observed in the assay will not always translate to what is observed *in vivo*, despite the strong concordance we observe with PRO-cap data.

However, the exogenous nature of SuRE also provides some advantages. By largely decoupling sequence activity from locus-dependent chromatin state, SuRE specifically assays DNA regulatory activity and can reveal regulatory potential that may be inactive in the specific assayed cell type but active in other cellular contexts. This potentially contributes to the observed SuRE signal across largely all promoters and broad enrichment of emVars among eQTL and GWAS signals linked to diverse tissues. Crucially, assaying the exact same DNA libraries in matched bovine and human endothelial cells enabled a highly controlled cross-species comparison of sequence effects, minimising confounding factors related to local genomic context and orthologous region structure. These features make SuRE particularly well-suited for dissecting conserved regulatory DNA grammar.

Motivated by the substantial scale of human functional genomics resources and sequence-based models, we directly tested the portability of DNA-encoded regulatory logic across species. Transfecting the cattle DNA libraries into matched primary cattle and human aortic endothelial cells revealed widespread regulatory conservation at multiple levels. Cattle regulatory elements largely retained autonomous activity in human cells. Likewise, allelic effects were predominantly directionally concordant across species, suggesting that many variants have qualitatively similar effects even if don’t reach significance thresholds due to differences in magnitude, power, transfection efficiencies, or cellular environments. This conservation extended to sequence modelling: promoter activity deep learning models trained independently on species-specific SuRE data learned broadly similar features, produced correlated predictions on the same cattle sequences, and showed comparable correspondence to observed SuRE activity. These results strongly support the view that a substantial fraction of regulatory “grammar”, particularly at promoters, is conserved across cattle and humans.

Despite this broad conservation, our cross-species design identified meaningful exceptions that highlight important considerations for transferring human regulatory annotations to livestock genomes. For example, we observed a divergence of regulatory effects at evolutionary young sequences and following the disruption of certain transcription factor motifs. Explicitly testing for species-by-allele interactions identified thousands of cross-species QTLs with significantly different allelic effects between cattle and human cells. Together, these findings support a pragmatic model for cross-species interpretation: while much of the baseline regulatory code is conserved and transferable, a non-trivial subset of sequences and variants violate simple transfer assumptions and will require explicit cross-species calibration.

In summary, by combining genome-wide SuRE profiling, graph-based haplotype-aware analysis, and direct cross-species assays in matched primary cells, we provide a high-resolution catalogue of cattle regulatory variants alongside a quantitative view of conserved regulatory grammar.

These resources and principles should accelerate causal variant discovery and trait improvement in cattle and offer a general framework for transferring, and appropriately fine-tuning, functional genomic knowledge across mammalian species.

## Methods

### Sample Acquisition and Cell Culture

The requirement for suitably transfectable primary cell lines, that could be grown to sufficient cell number for SuRE and where the cell physiology broadly matched between humans and cattle led to the focus on aortic endothelial cells. The primary human aortic endothelial cell line (HAEC-D1) was obtained from Lonza and cells were cultured in EBM-2 Basal Medium (Lonza, CC-3156) supplemented with EGM-2 SingleQuots (Lonza, CC-4176). The primary bovine aortic endothelial *B. taurus* cell lines (BAOEC-D1 and BAOEC-D2) were obtained from Creative Bioarray (CSC-C4393X) and were cultured in DMEM supplemented with 10% FBS, 1% GlutaMAX, and 1% Sodium Pyruvate Solution. Both HAEC-D1 and BAOEC cell lines were passaged using 0.25% trypsin (Westburg, CA TRY-3B).

*B. indicus* cells (Bi-83-A) used to obtain DNA for the SuRE library were previously derived from a Sahiwal calf and immortalised via infection with *Theileria annulata*^32^. These suspension cells were cultured in RPMI-1640 medium supplemented with 10% FBS, 1% GlutaMAX, 50 µM beta-mercaptoethanol, and penicillin/streptomycin. They were passaged by scraping the flask to release loosely adhered cells and transferring one-quarter of the volume to fresh medium. All cell cultures were screened for mycoplasma every 3 months using PCR (Takara, #6601).

### Genome Graph Construction and Genotyping

Genomic DNA samples were prepared from cultured Bi-83-A and BAOEC-D1 cells and sequenced on the PacBio Revio platform at Edinburgh Genomics. Dual genome assemblies were then generated using hifiasm^33^. Read counts, effective coverages and assembly statistics are shown in Supplementary Table 4. A genome graph was generated using Minigraph-Cactus^34^ from these four assemblies, alongside the bovine reference genome, ARS-UCD2.0^17^, and the *T. annulata* reference genome ASM322v1^35^ (due to the presence of *T. annulata* DNA in the *B. indicus* SuRE libraries). To generate a set of variants for analysis, the VCF generated by Minigraph-Cactus was filtered to remove variants with missing genotype calls or where multiple alleles were not observed at least once across the two source cattle samples.

### SuRE MPRA Library Preparation and Sequencing

SuRE libraries were generated as previously described^13^. Cattle DNA for both samples were isolated using the DNeasy Blood & Tissue Kit (Qiagen, 69504). DNA was fragmented as previously described^13^ and size-fractionated using CleanNGS beads (CLEAN NA, CNGS-0050) to isolate elements in the ∼400–600 bp target range. Libraries were generated for each genome via transformation into E. cloni 10G (Lucigen, 60107-1).

For the human cell assay, 45 µg of the mixed library was transfected into 50 confluent 15 cm plates of HAEC-D1 cells per technical replicate using ViaFect Transfection Reagent (Promega, E4982) according to the manufacturer’s instructions. The bovine assay was performed identically using BAOEC-D1 and BAOEC-D2 cells, except 30 confluent 15 cm plates were transfected per biological replicate.

RNA was isolated directly on the plates using TRIzol (Fisher Scientific, 12034977). For each replicate, 3 mg of RNA was processed into cDNA in a 12 ml reaction volume, which was subsequently PCR-amplified in a 60 ml total volume. A 200 µl aliquot of this reaction was purified using CleanNGS beads and sent for Illumina short-read sequencing. Both the initial input plasmid library (”input”) and the cDNA-derived libraries (”cDNA”) were sequenced. Total read counts are shown in Supplementary Table 5.

### Fragment Data Processing

Barcodes were isolated and enumerated from the cDNA sequencing data to quantify the number of transcripts derived from the DNA fragment associated with each barcode. To identify the genomic position of each library fragment, input sequencing reads were split into barcode and fragment sequences.

For analyses based on a linear reference genome (peak calling, comparison to non-graph based emVar calling and sequence-to-transcription modelling) fragment sequences were aligned using a haplotype-aware workflow. Briefly, two pseudo-haplotype-specific genome sequences were generated for each cattle subspecies using the bcftools^36^ consensus function. The input VCFs were derived from Illumina short-read WGS data called relative to the ARS-UCD1.2 reference genome. Trimmed, paired-end, reads were aligned to the reference genome using the BWA-MEM algorithm within BWA-Kit^37^, which additionally marks duplicates and adds read-group information. Median coverage was >21x for both individuals. Per-sample variants were called using DeepVariant v1.4.0^38^, and multi-sample, bi-allelic, joint-genotyped calls with GLnexus^39^. Missing genotypes were imputed, and the dataset phased, against a reference panel from the 1000 Bulls Genomes Project^40^ (Run 8) with Beagle v.5.4^41^.

Input sequencing reads were then aligned to both haplotype-specific reference sequences using Bowtie2^42^. Resulting alignments were classified as haplotype-specific, identical between haplotypes, or ambiguous. Ambiguously aligned fragments were discarded. Fragments with unambiguous alignments were subsequently lifted back to standard reference genome coordinates using the UCSC liftOver tool^43^. Finally, these alignments were matched to cDNA counts via fragment barcodes to generate tab-delimited files containing fragment occurrences in the input library, corresponding cDNA counts across species and replicates, and genome-wide SuRE signal tracks.

For graph-based variant analysis, fragment sequences were aligned to the genome graph using Giraffe^44^. Aligned reads were processed to associate them with their barcode and cDNA counts, and to aggregate reads originating from the same fragment and fragments with identical start and end locations. Quality control included removing reads with invalid barcodes and removing fragments associated with barcodes that appeared in multiple reads mapped to different positions.

To normalise SuRE scores for the fragments, a normalisation factor was calculated for each library by dividing the total cDNA counts for that library by its total iPCR count for all fragments. A fragment SuRE score was then calculated by dividing the fragment cDNA counts by its iPCR count, and dividing this by the appropriate normalisation factor. To produce the fragment plots, the start and end of the graph-based fragment alignments in the reference genome were calculated based on the first node in the read alignment that was present in the reference path.

To enable identification of the variants and alleles carried by each fragment, the paths of each allele in the graph provided by the Minigraph-Cactus VCF (“AF” INFO field) were parsed to identify the minimum path unique to each allele. For example, for a deletion variant with alleles “>99>100>101” and “>99>101” (denoting a deletion of node 100 between shared flanking nodes 99 and 101), these minimum paths would be “>100” for the reference allele and “>99>101” for the alternative allele. For SNVs, the flanking nodes were added to these paths to ensure the single differing base was evaluated in the expected sequence context. The read alignment paths output by Giraffe were then searched for these unique allele paths. When paired reads for a fragment did not overlap, alignment paths were compared with the sample’s haplotype paths in the graph. If these matched exactly one haplotype, genotypes of variants from the unsequenced region between the reads were inferred from the haplotype path; if reads matched both haplotypes, only homozygous variant genotypes were inferred. The entire analysis from input sequencing to final count files was orchestrated using a reproducible Nextflow workflow (see code availability section).

### Peak Calling and Annotation

SuRE expression scores are defined as the ratio of cDNA to iPCR read counts at each genomic position, scaled such that the genome-wide mean is 1. To identify regions of high transcriptional activity, peaks were called from these SuRE expression signals using the peak-calling tool PINTs^45^ (Peak Identifier for Non-canonical Transcripts). BigWig files from both replicates on the forward and reverse strands were used as inputs to pints_caller for peak calling, which generated both bidirectional and unidirectional peaks. To assign strand orientation to bidirectional peaks, we identified the nearest transcription start site (TSS) for each peak using the BEDTools^46^ closest function and assigned the strand of the nearest TSS to the corresponding bidirectional peak. Bidirectional and unidirectional peaks were subsequently merged, and only peaks passing the PINTs q-value significance threshold (0.05) were retained for downstream analyses.

Genomic annotations were derived from the *B. taurus* ARS-UCD2.0 Ensembl GFF3 file (release 114). Introns were defined as transcript regions excluding exons, promoters were defined as the region 10kb upstream of TSSs in a strand-aware manner, and intergenic regions were defined as the complement of merged genic intervals. SuRE peaks were annotated by intersecting peak coordinates with genomic features using BEDTools^46^, and each peak was assigned to a single category using a hierarchical priority (promoter, 5’ UTR, 3’ UTR, exonic, intronic and intergenic).

### Epigenomic Profiling

Cattle heart ChIP-seq data for H3K4me3 and H3K27ac were obtained^47^ (sample accessions SAMEA4447809, SAMEA4675151 and SAMEA4447801). Raw sequencing data were processed using the nf-core/chipseq^48^ pipeline with default parameters to generate aligned BAM files. Chromatin enrichment at SuRE peaks was quantified using HOMER^22^ (annotatePeaks.pl). SuRE peaks were first classified as proximal (≤10 kb from the nearest TSS) or distal (>10 kb). ChIP-seq signal for H3K4me3 and H3K27ac was quantified for each peak using HOMER tag directories generated from the aligned BAM files, summarising read densities within a 5kb window centred on each peak.

### PRO-cap Data Generation and Processing

PRO-cap libraries were generated for the cattle aortic endothelial cells by adapting the protocol of Judd et al.^49^ for use in bovine cells, with all reactions carried out in the presence of RNase inhibitor. Briefly, 5x10^6^ cells were washed in Dulbecco’s Phosphate Buffered Saline (DPBS), permeabilized in ice-cold permeabilization buffer, and washed twice in wash buffer before being resuspended in reaction buffer for the nuclear run-on reaction. The permeabilized cells were combined 1:1 with 2X Run-on Master Mix containing 40 µM Biotin-11-UTP and 40 µM Biotin-11-CTP, and incubated at 37°C for 5 minutes. Following the run-on reaction, RNA was extracted using TRIzol, cleaned up using Bio-Rad Micro Bio-Spin P-30 Gel Columns, and precipitated with ethanol and GlycoBlue. The resulting RNA pellet was resuspended in 1.5 µM of Illumina TruSeq Small RNA Adapter RA3 containing a 6 bp UMI, denatured at 65°C, and subjected to 3’ adapter ligation using T4 RNA Ligase 1 at 25°C for 1 hour.

Biotinylated transcripts were then enriched using Dynabeads MyOne Streptavidin C1 beads. Prepared beads were added to the ligated RNA samples and incubated with rotation for 20 minutes at room temperature. The beads were sequentially washed with high-salt and low-salt buffers, followed by on-bead 5’ de-phosphorylation using Antarctic Phosphatase for 30 minutes at 37°C, and 5’ hydroxyl repair using RNA 5’ Pyrophosphohydrolase (Rpph) for 1 hour at 37°C. After removal of the supernatant, the beads were resuspended in 1.5 µM of Illumina TruSeq Small RNA Adapter RA5 with a 6 bp UMI, denatured at 65°C, and subjected to 5’ adapter ligation using T4 RNA Ligase 1 at 25°C for 1 hour. The beads were washed again with high-salt and low-salt buffers, and RNA was subsequently eluted from the beads using TRIzol and precipitated with ethanol and GlycoBlue.

The recovered RNA was resuspended in RT resuspension mix, denatured at 65°C, and reverse-transcribed using SuperScript III Reverse Transcriptase at 50°C for 45 minutes, followed by enzyme denaturation at 75°C. For library amplification, unique Illumina TruSeq Small RNA RPI-n index primers were added to each sample alongside Q5 Hot Start High-Fidelity DNA Polymerase and amplified using standard thermocycling conditions. The resulting PCR products were cleaned using AMPure XP beads at a 1:1.8 ratio and eluted in 10 mM Tris-HCl (pH 8.0). Finally, library quality and fragment size were confirmed using a D1000 TapeStation, and libraries were quantified via a Qubit dsDNA Assay prior to pooling and sequencing on an Illumina NextSeq 2000 P3 platform (2x 100 bp).

Multiple libraries were sequenced on both Illumina MiSeq and HiSeq platforms, and reads from each run were processed independently using the same pipeline. In total 1,477,561,133 raw read pairs were obtained across all sequencing runs. Sequencing reads were processed using fastp^50^ to remove adapter sequences, trim low-quality bases, and discard reads shorter than 14 bp. Unique molecular identifiers present at the start of both reads were extracted using UMI-tools^51^ and appended to read headers. Processed reads were aligned to the reference genome using STAR^52^, generating coordinate-sorted BAM files. BAM files from different runs were subsequently merged using samtools^53^. PCR duplicates were removed using UMI-tools in paired-end mode based on the extracted UMIs, with deduplication performed either before or after merging depending on the dataset. The final merged and deduplicated BAM file was indexed using samtools for downstream analyses, yielding 3,748,076 read pairs.

To validate SuRE activity against endogenous transcriptional initiation, we compared SuRE expression scores with PRO-cap signals. PRO-cap alignments were processed using bamCoverage^54^ to generate signal tracks at 1-bp resolution. Alignments were filtered for a minimum mapping quality (MAPQ) of 20, and PCR duplicates were removed to ensure high-fidelity signal representation. The resulting 5’ end nascent transcript signals were normalized to Counts Per Million (CPM) and exported as BigWig files. For comparison, mean signals from both assays were extracted within 2 kb windows centred on the transcription start sites of annotated *B. taurus* protein-coding genes. To assess the relationship between reporter activity and endogenous initiation, Pearson (r) correlation coefficients were calculated on log_10_-transformed mean values. To minimize the impact of stochastic noise in low-signal regions, the analysis was restricted to TSS windows exceeding the 5^th^ percentile of signal intensity in both datasets.

### Variant Filtering and Allele-Specific Expression

Biallelic variants were filtered using thresholds calculated for each combination of library (*B. taurus, B. indicus*) and genome graph genotype calls (0/0, 0/1, 1/1). The mean and standard deviation of the proportion of fragments carrying the reference allele were calculated for each combination, considering only fragments where the allele was determined directly from the sequence. Variants falling outside the mean +/- 2.58 standard deviations for their corresponding genotype were discarded as this was suggestive of genotype call discrepancies between the graph and SuRE data. To exclude sites with unusual coverage, variants were discarded if their total fragment occurrences exceeded 1000 or fell outside +/- 2.58 standard deviations from the mean of remaining variants. Lastly, variants were discarded if the proportion of fragments where the allele was inferred from haplotype paths rather than observed from aligned sequence reads was greater than 2.58 standard deviations from the mean, as this indicated that the genotype was not well supported by sequencing data. At the remaining sites, any fragments that carried alleles inconsistent with the genome graph genotypes were removed. Since filtering was first carried out at the library level, if the variant’s fragments from one library had fallen outside the thresholds described, the variant would remain in the analysis using only fragments from the second library, as long as these carried both reference and alternative alleles. 3.7% of variants were filtered out (n=639,786), and 1.9% of retained variants had spanning fragments from a single library only (n=322,494).

Filtering of variants which were not biallelic in the genome graph was performed in an analogous way, using the thresholds calculated from the biallelic data, using the 1/1 thresholds for any non-reference homozygotes and the 0/1 thresholds for any heterozygotes. An additional filter was added to remove any variants where the proportion of fragments carrying an allele inconsistent with the graph genome genotype was outside the mean +/- 2.58 standard deviations for all non-biallelic variants. 11.4% of variants were discarded from both libraries, and 13.8% had fragments from only one library remaining after filtering, an increase over the proportion for biallelic variants that suggested these were more likely to be the result of variant calling errors. These variants (n=79,705) were therefore also discarded from the analysis. For the retained variants, if only two alleles were segregating in the SuRE fragments (e.g. genotypes of 1/1 and 2/2, n=126,539), these were recoded to alleles 0 and 1 and tested using the same method as the biallelic variants (see “Modelling of emVars” below). However, because these variants no longer necessarily matched the original VCF in alleles or position, they were excluded from some analyses. They were included in the counts for total and regulatory variants in cattle cells, the corresponding plots in Figures 2a and 3a, and in analyses of the effect of variant size, but no other analyses. The remaining, filtered multiallelic variants (n=305,466) had at least three alleles represented on SuRE fragments.

### Modelling of emVars

To model the regulatory impact of biallelic DNA variants, we utilized zero-inflated negative binomial (ZINB) regression. This accommodated over-dispersion of cDNA counts and an excess of zero counts arising from stochastic plasmid loss during transfection. For each DNA fragment *i*, the total cDNA count *Y*_*i*_ was modelled as the dependent variable. We included the logarithm of the corresponding input DNA count as an offset to normalise for differences in fragment abundance.

The negative binomial component estimating the regulatory activity of transcribed fragments was defined as:

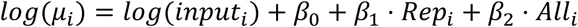

Where *input*_*i*_ is the fragment count, *Rep*_*i*_ indicates the replicate, *All*_*i*_ is an indicator for the allele (0 = reference, 1 = alternate) carried by the fragment, and *β*_0_,*β*_1_and *β*_2_ are estimated coefficients.

The zero-inflation component was modelled as:

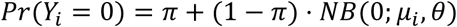

Where *π* is the estimated zero-inflation probability, and *θ* is the dispersion parameter.

To disentangle species-specific effects, analyses were conducted separately for fragments assayed in cattle cells and human cells. Sites showing a statistically significant effect of the allele after Benjamini-Hochberg correction were classified as putative regulatory variants (emVars). In some cases (n = 4,893 and n = 6,955 for transfections into cattle and human cells, respectively) the model failed to converge. For the cross-species analyses we discarded any variants where this occurred for data from either species. Variants where the normalised fragment SuRE score values had a skew of >20 were processed, but not included in the final data set (n = 84,599 for high skew in cattle cell data (excluded from all analyses), n = 35,582 for high skew in human cell data (only excluded in comparisons of transfections in cow and human cells).

For non-biallelic variants a similar approach was used, but the allele carried by the fragment could no longer be expressed as an indicator variable since there were more than two possibilities. Instead, a model including the allele as a covariate (model 1) was compared with a simpler model that did not include the allele (model 2). Model 1 was identical to the biallelic model above except that *All*_*i*_ indicates the allele carried by the fragment encoded as a factor. In model 2 the zero-inflation component was identical and the negative binomial component was defined as:

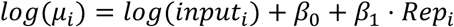

Where *input*_*i*_ is the fragment count , *Rep*_*i*_ indicates the replicate, and *β*_0_ and *β*_1_ are estimated coefficients. The fits of the two models were compared using a likelihood ratio test using R, and the P-values adjusted using the Benjamini-Hochberg procedure. Since multi-allelic variants were tested using a different model from the biallelic variants and their individual allele effects were not determined, the P-values of the two variant types were not comparable, and the multi-allelic variants were not included in genome-wide analyses. Variant filtering and modelling was carried out using a reproducible Nextflow workflow (see code availability section).

All analyses were performed in R 4.4 utilizing packages including pscl^55^ for zero-inflated models and future/furrr^56^ for parallelisation.

### Population Genetics and Allele Frequencies

To investigate the link between regulatory activity and population divergence, we compiled public whole-genome sequencing data from 56 taurine and 56 indicine samples (Supplementary Table 6). Genotyping was performed using PanGenie^57^ v4.2.1 against the genome graph, and allele frequencies were calculated using bcftools^36^ v1.6. Derived Allele Frequency (DAF) was annotated using ancestral allele information previously derived from a multi-species alignment^58^.

We performed logistic regression to model whether an SNV was a cattle emVar. The probability that variant *ii* was regulatory (p*i*_*i*_) was modelled as:

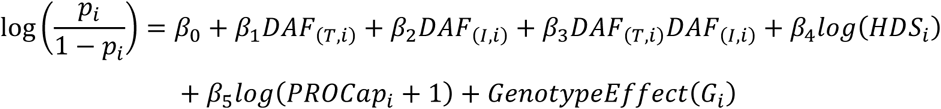

Where *DAF*_(*T*,*i*)_ and *DAF*_(*I*,*i*)_ denote the variant’s taurine and indicine derived allele frequencies respectively, *HDS*_*i*_ is the variant density score within a 1kb window centred on variant *i*, *P*ROCap_*i*_ is the maximum PRO-cap read count within 500bp and categorical genotypes of the source animals were included to control for their influence.

### Conditional Analysis of Local Haplotypes

To evaluate within breed linkage disequilibrium among the regulatory variants we utilised previously generated variant calls^59^ from a cohort of 88 high coverage Holstein-Friesian samples (Bioproject ID: PRJEB27379). Relatedness2 in vcftools^60^ was run with a MAF cutoff of 0.05 and max-missing of 0.8 to identify related samples. Samples were then iteratively excluded until none left had a score > 0.0625 with another remaining sample, leaving the final set of 33 unrelated samples. Variants were restricted to biallelic SNPs genotyped in this population and with a missing call rate less than twelve samples, leaving 123,970 variants. Pairwise LD (r^2^) calculations were performed using PLINK^61^ (v1.90b). Calculations were restricted to variant pairs separated by a physical distance between 5 kilobases and 1 megabase.

To search for variants that are close together but have separate effects, pairs of biallelic regulatory variants within 200 bp of one another and carried together on at least 10 fragments were identified. Where clusters of multiple sites were linked together in successive pairs, the emVar with the lowest P-value was taken as the focal variant and only pairs containing this variant were tested. From this list, variant pairs in complete linkage were then discarded. Many fragments carried other variants in addition to emVars; most of these had a non-significant result in the test for biallelic allele-specific effects and were assumed to have no effect, but for a range of reasons some variants could not be tested (e.g. missing genotypes, non-biallelic variants, filtered out as described in the Variant Filtering and Allele-Specific Expression section). To avoid confounding effects from these, any variant pair with such an untestable variant on more than half of its fragments was discarded from the analysis.

To test for independent effects in the remaining pairs, the biallelic allele-specific effect model described above was refitted for the focal variant using only fragments spanning both variant. Its fit was then compared with that of a second model, identical except that negative binomial component was defined as:

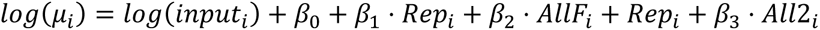

Where *AllF*_*i*_ is an indicator for the allele carried by the fragment at the focal variant, and *All*2_*i*_ is an indicator for the allele carried by the fragment at the second variant (0 = reference, 1 = alternative). A significantly better fit of the second model was determined by a likelihood ratio test using R, and the P-values adjusted using the Benjamini-Hochberg procedure. An FDR ≤0.05 was interpreted as evidence both variants having potential independent regulatory effects.

### eQTL and GWAS Enrichment Analysis

To validate the *in vivo* relevance of our MPRA findings, we tested for an enrichment of regulatory variants within the cattle GTEx^19^ (cGTEx) project expression quantitative trait loci. We processed the eQTL data to retain only the single most significant gene association for any SNP linked to multiple genes. The SuRE SNVs were merged with these filtered eQTLs by SNP coordinates and partitioned into four groups: (i) significant emVar in both cattle and human cells, (ii) significant in human cells only, (iii) significant in cattle cells only, and (iv) not significant in either. A one-sided, two-sample Wilcoxon rank-sum test was used to determine if the distribution of cattle GTEx eQTL P-values for each of the three significant emVar groups was stochastically lower than the P-value distribution of the ’not significant’ background group. Pairwise LD values for the visualisation in Figure 2d were calculated using the Holstein animals from the 1000 Bull Genomes Project^40^ using PLINK^61^.

To assess enrichment of SuRE regulatory variants near cattle trait loci, we focused on fine-mapped variants associated with diary production traits obtained from published cattle GWAS fine-mapping analyses. We obtained 4,296 fine-mapped variants (3,498 after removing duplicates) across six traits, including milk yield, milk fat yield, milk protein yield, milk fat percentage, milk protein percentage and somatic cell score^62–64^. For comparison, background SNPs were generated by matching each fine-mapped variant by chromosome and minor allele frequency (MAF). MAF was calculated using taurine allele frequencies (see Population Genetics and Allele Frequencies), reflecting the predominance of taurine cattle in the fine-mapped population. For both fine-mapped and matched background variants, the genomic distance to the nearest SuRE regulatory variant was calculated and grouped into distance bins to compare the distribution of distances between fine-mapped variants and matched controls.

### Testing for Species-Specific Effects (xs-emVars)

To identify cross-species emVars (xs-emVars) exhibiting significantly different effects between cattle and human cells, we conducted a comparative analysis of the estimated allele effects (*β*_2_) obtained from the ZINB models. For each variant, we computed a Wald test statistic:

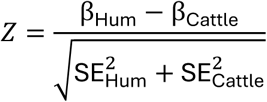

Where β and *SE* represent the estimated variant effect sizes and their corresponding standard errors in human and cattle cells. A two-sided P-value was calculated and adjusted using the Benjamini-Hochberg procedure to control the false discovery rate (FDR).

### Transcription Factor Motif Disruption Analysis

To investigate potential transcription factor (TF) mechanisms underlying xs-emVar effects, we evaluated whether xs-emVar variants altered predicted TF binding motifs using motifbreakR^65^. Variants were supplied as genomic coordinates (BED) and analysed using motifbreakR with position weight matrices (PWMs) from bundled motif collections, including HOCOMOCO, HOMER, FactorBook and ENCODE motif databases. Motif disruption was assessed using the information-content scoring framework with a significance threshold of 1×10⁻⁴, retaining strong, weak and neutral calls. To reduce redundancy from multiple PWMs corresponding to the same TF, motif hits were collapsed to retain the strongest predicted disruption per TF for each variant based on the largest absolute allele-specific score difference.

To test whether TF motifs were preferentially disrupted among xs-emVars, we compared motif hit frequencies between xs-emVars and background variants defined as cattle emVars not also classified as xs-emVars. Variants with at least one motif disruption passing an allele-effect filter (|alleleEffectSize| ≥ 0.15) were counted for each TF, retaining the strongest hit per variant-TF pair. Enrichment was evaluated using two-sided Fisher’s exact tests with P values adjusted using Benjamini-Hochberg procedure.

### Sequence-to-Function Model (EvaReg)

We extracted the locations of 35,351 protein-coding TSS regions (autosomes only) from Ensembl (Ensembl release 115). Sequence blocks were defined by extracting 300 bp upstream and downstream of the TSS. Data augmentation was performed by applying strides, shifting the block centre by 100 bp, consecutively 3 times on either side, resulting in 7 sequence blocks per TSS. Reverse-complemented sequences were also included for robustness.

To calculate a continuous SuRE score for training, we aggregated fragments overlapping the sequence block. A weighted contribution score was computed as:

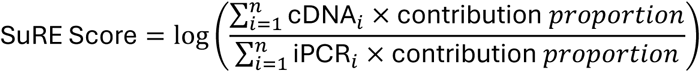

Where we have *ii* fragments overlapping a given sequence block. Where the contribution proportion is the fraction of base pairs overlapping the sequence block window. Since some resulting scores were negative, we shifted all scores up to ensure a minimum value of 0 by adding the absolute value of the smallest value.

Inspired by InceptionNet^66^, EvaReg takes 600 bp inputs to predict regulatory activity. The architecture begins with 3 convolution blocks of varying kernel sizes (7, 1, and 3) to capture broad and specific motif features. This is followed by 9 consecutive Inception blocks, which apply parallel operations with different receptive fields before concatenation, utilising AttentionPool layers to select critical regions. The network concludes with an AdaptivePool layer, a Dropout layer (p=0.4), and a Linear layer for regression output.

Chromosomes 8 and 13 were held out for final testing. Hyperparameters were optimized via Optuna^67^ over 30 trials using a 3-fold cross-validation scheme (30 epochs per fold). Distributed training was conducted across 6 NVIDIA H100 GPUs.

### Variant Effect Model Prediction

To test model performance at variant annotation, SNPs located within 300 bp of a TSS were categorised into ’significant’ (adjusted p < 0.001) and ’not significant’ (adjusted p > 0.8) based on the cattle emVar data, generating 405 significant SNPs and 41,417 non-significant SNPs across the whole promoter sequence dataset. 600-bp sequences, centred around the TSS and containing the variant, with the reference and alternate alleles, were passed through the model. The absolute change between reference and alternate predicted scores was evaluated using the Kolmogorov–Smirnov test. For significant SNPs, the direction of predicted change was compared to experimental beta values using a binomial test.

### Motif Activity throughout the Cattle Genome

For the baseline model importance score throughout the entire dataset, for each promoter, the importance score was defined as the difference between the reference base at a given position and the mean of the alternate bases. The score at each base was further divided by the SuRE activity score of the given promoter’s reference sequence to account for the difference in activity levels across different sequences. The baseline score was further z-score normalised relative to the entire dataset.

For each motif instance, the score was extracted by taking the average of the importance scores across the bases it spanned. The motif instance was labelled as such:

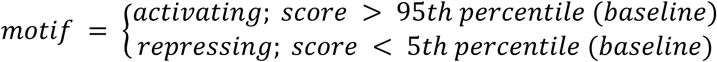

Furthermore, we have noted down the location of occurrence of such ‘activating’ or ‘repressing’ motifs from their respective TSSs to observe the distribution difference between activating and repressing motifs.

### Motif Analysis Across Species

To characterise the motif activity conservation across species, we extracted the importance scores corresponding to each motif from the baseline score for the entire promoter dataset. For the distribution of importance scores corresponding to a motif in either species, we compared them against their corresponding species baseline score using the Kolmogorov–Smirnov test. The observed P values were adjusted using the Benjamini-Hochberg procedure. For Figure 5f the minimum of the resulting q-values was taken across the cattle-in-cattle and cattle-in-human analyses. For motif instances with a q-value of 0, scores were replaced with the next lowest q-value in the whole promoter dataset.

### Conservation, Repeats, Motif Enrichment and Orthology Mapping

The evolutionary conservation of regulatory regions was assessed using cattle phyloP conservation scores^68^, which were previously calculated based on multiple sequence alignments of 241 vertebrate species. Peaks were grouped into cattle-specific, human specific, and shared regulatory peaks, and conservation profiles were computed using deepTools^54^. PhyloP scores were extracted in reference-point mode centred on the peak midpoint using computeMatrix, within a symmetric window (10 kb) around the peak centre and summarized in 10-bp bins. Positions lacking phyloP signal were excluded. To evaluate whether the reduced conservation observed at human-specific peak centres was driven by repetitive elements, RepeatMasker^69^ annotations for the cattle genome (ARS-UCD2.0) were obtained from the UCSC Genome Browser^70^, peaks overlapping annotated repeats were removed, and conservation profiles were recomputed using the same workflow.

To identify orthologous regions in the human^71^ (T2T-CHM13) and pig^72^ (T2T-pig1.0) telomere-to-telomere genomes around emVars we performed targeted cross-species sequence alignments. Single nucleotide coordinates of emVars mapped to the cattle reference genome^17^ (ARS-UCD1.2) were symmetrically expanded by 25 base pairs using BEDTools^46^ (v2.31.1) to generate 51-bp query windows. Nucleotide sequences for these regions were then extracted from the cattle reference using the BEDTools getfasta utility.

The resulting cattle query sequences were aligned to the human or pig reference genome assemblies using LASTZ^73^. To optimize for cross-species homology identification while maintaining high mapping specificity, alignments were executed using the following parameters: --step=1, --seed=12of19, --notransition, and --hspthresh=1800.

Following alignment, results were parsed and filtered using R. To ensure each cattle locus was represented by its most likely ortholog in each species, redundant alignments were filtered, retaining only the single highest-scoring alignment for each unique cattle query.

## Acknowledgements

This work was primarily supported via funding from the UK’s Biological and Biotechnology Research Council (BB/W000288/1, BBS/E/RL/230001B). This project was also partly supported via funding from the Gates Foundation (INV-076519) and with UK aid from the UK Government’s Department for International Development (Grant Agreement OPP1127286) under the auspices of the Centre for Tropical Livestock Genetics and Health (CTLGH), established jointly by the University of Edinburgh, SRUC (Scotland’s Rural College), and the International Livestock Research Institute (ILRI). The findings and conclusions are those of the authors and do not necessarily reflect the positions or policies of the BMGF or the UK Government. We thank Miriam Smits for generating the SuRE libraries and Michelle van Heck for performing tissue culture and transfection experiments. We also acknowledge funding from a Roslin Foundation grant (RF2301). We acknowledge the use of the EIDF GPU Service, a service operated by EPCC at the University of Edinburgh’s Advanced Computing Facility and funded by the DDI programme of the Edinburgh and South-East Scotland City Region Deal.

## Data availability

All relevant data, including raw sequencing files, will be made available upon acceptance.

## Code availability

The code related to the methods and results in the paper are available on GitHub at: https://github.com/evotools/SuRE-paper-codes.git

