## Supplementary Figures 1-12 and Tables 1-6 for "A high-resolution atlas of cattle regulatory variants and their cross-species activity in matched human cells"

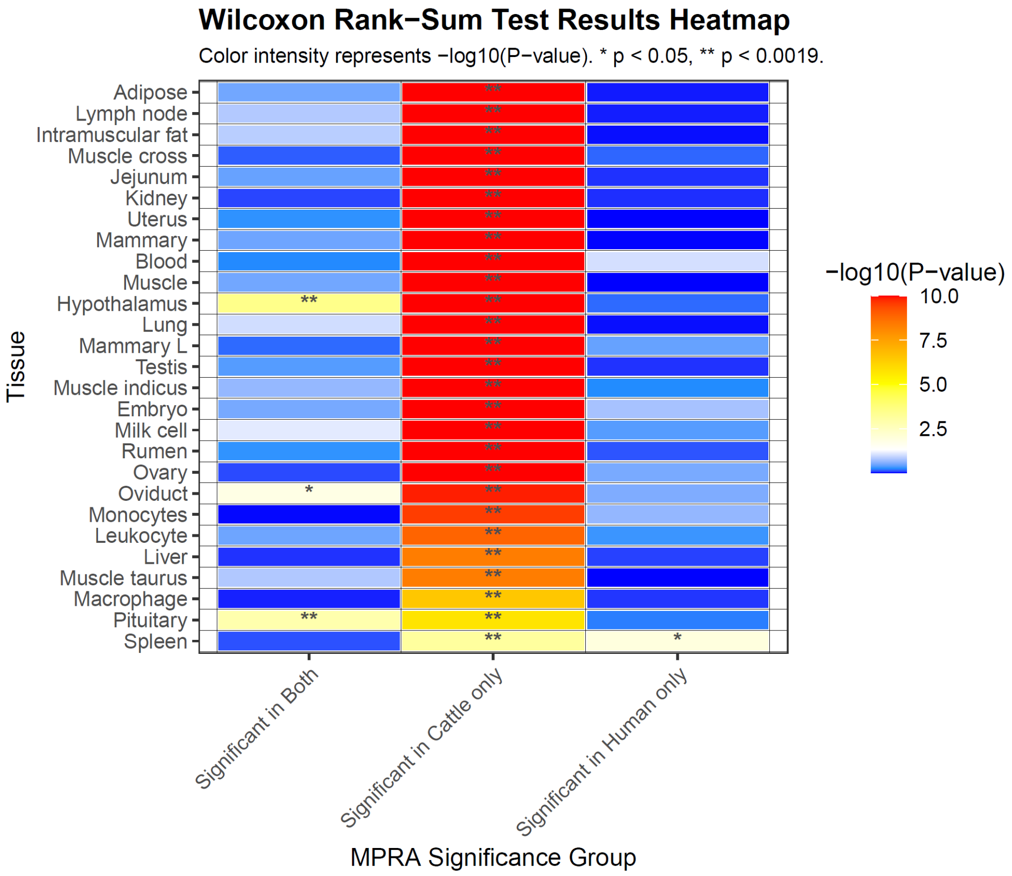


**Supplementary Figure 1.** Enrichment of cattle GTEx eQTL signal among MPRA-significant variants, by tissue (Wilcoxon tests). Heatmap summarizes two-sample Wilcoxon rank-sum tests performed separately for each cattle GTEx tissue (rows) and each emVar class (columns). It can be seen that emVars identified in cattle cells are significantly enriched at eQTLs identified across the range of cell types (middle column). emVars significant in both human and cattle cells, or human cells only, that are described later, are included for comparison. For each variant we used the minimum nominal cattle GTEx P-value across genes for that SNP in that tissue. Cell colour shows $-\log_{10}(P)$ for the Wilcoxon test of whether $-\log_{10}$(cattle GTEx P) is stochastically larger in the SuRE-significant group than in the non-significant group (values above 10 are shown as 10). * $P < 0.05$; ** $P < 0.0019$. Tissues are ordered by the P-value in the cattle-only emVar column.

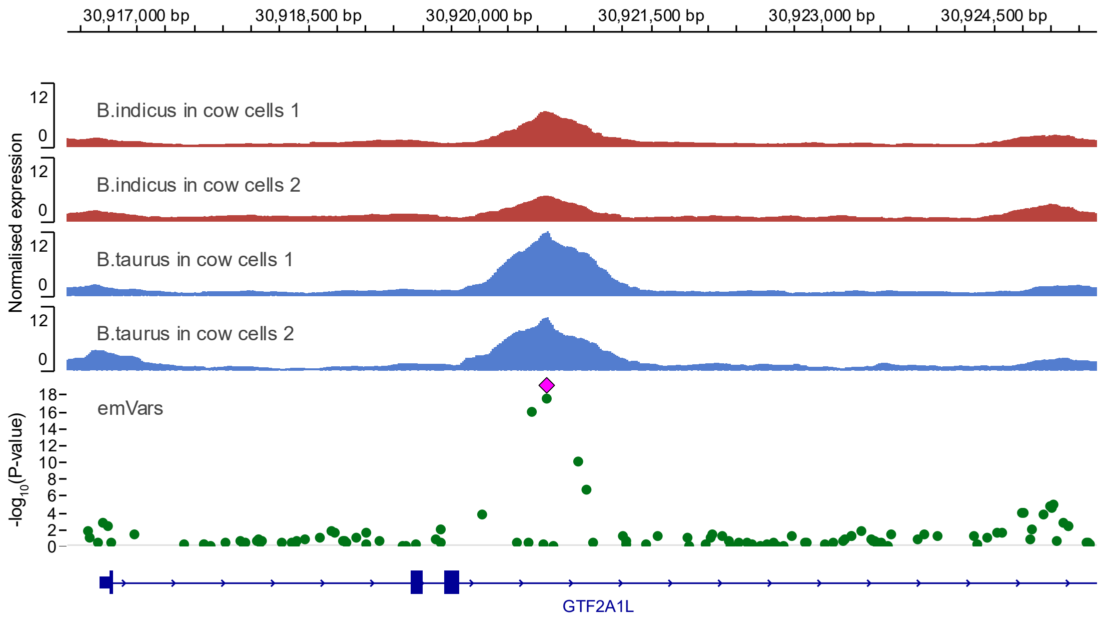


**Supplementary Figure 2**. SuRE (red and blue), and emVar (green points) signal tracks around the locus shown in Figure 3 panel (b), in exon 3 of the *GTF2A1L* gene. The point representing the 11:30920596 insertion is indicated in magenta.

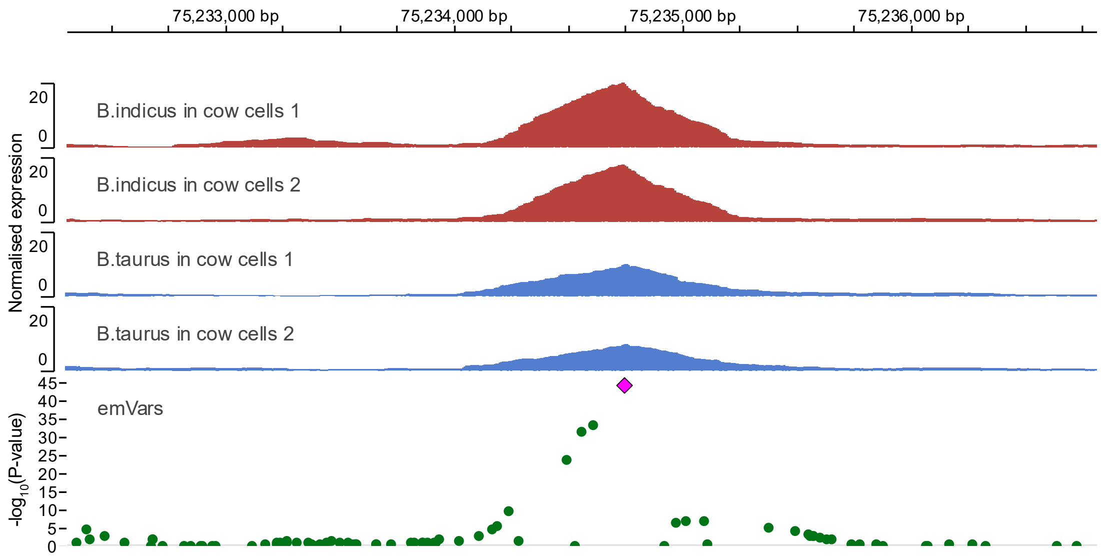


**Supplementary Figure 3.** SuRE (red and blue), and emVar (green points) signal tracks around the locus shown in Figure 3 panel (e), 21 kb upstream of lncRNA gene ENSBTAG00000062845. The point representing the multiallelic variant at 5:75234745 is indicated in magenta.

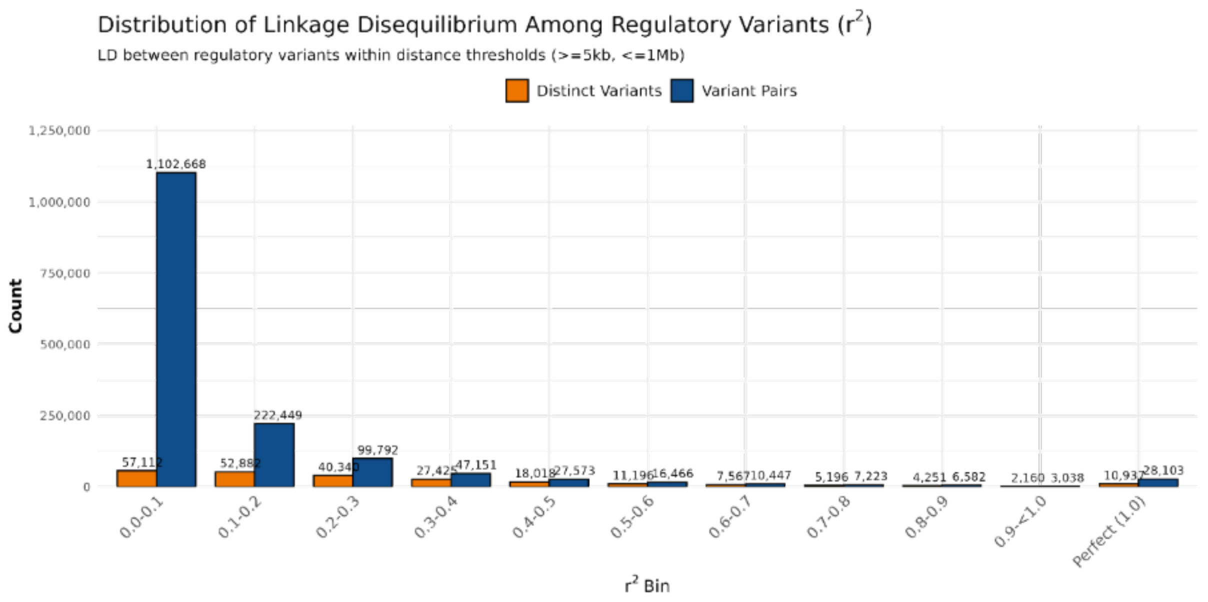


**Supplementary Figure 4**. Linkage disequilibrium observed among pairs of regulatory variants in Holstein-Friesian cattle. Pairwise LD was calculated for significant emVar variants separated by at least 5 kb and less than 1 Mb across all 29 autosomes. The x-axis categorises $r^2$ values into decile bins, including a distinct bin for variant pairs in perfect LD. The blue bars denote the absolute count of variant pairs falling into each bin. The orange bars indicate the number of distinct (unique) individual variants involved in those pairs, illustrating the extent to which single variants participate in multiple LD pairings.

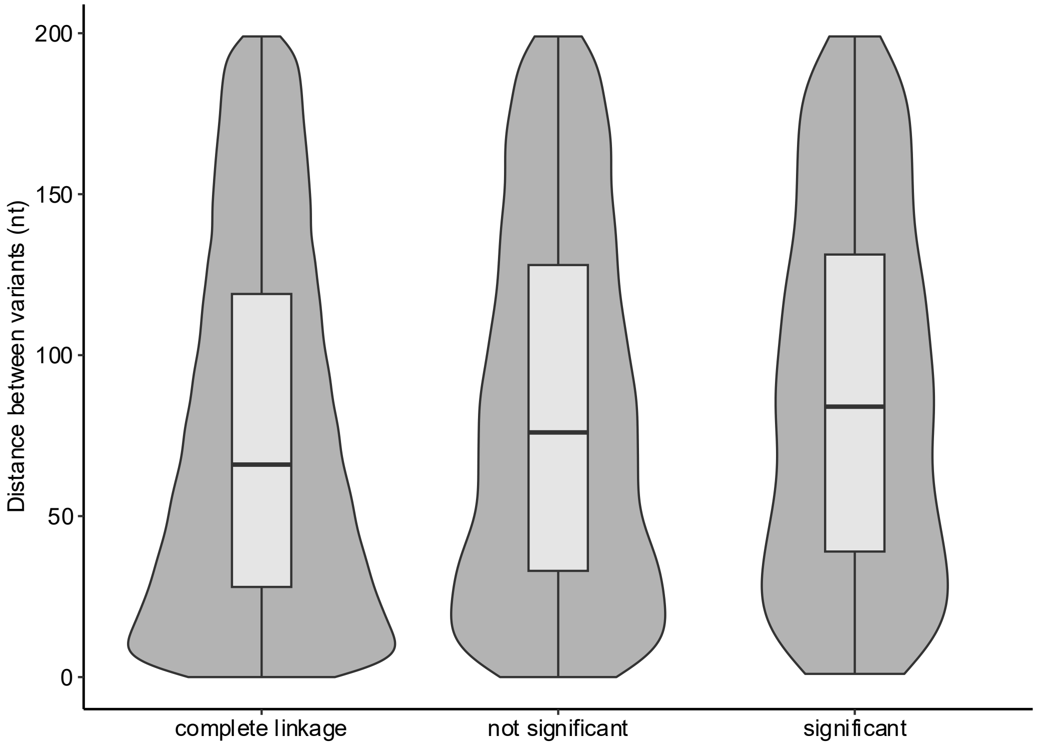


**Supplementary Figure 5**. Combined violin and box plot of pairs of variants frequently found on the same fragment, classed by whether they were in complete linkage, had no significant additional effect of the second variant, or had evidence of independent regulatory effects ("significant").

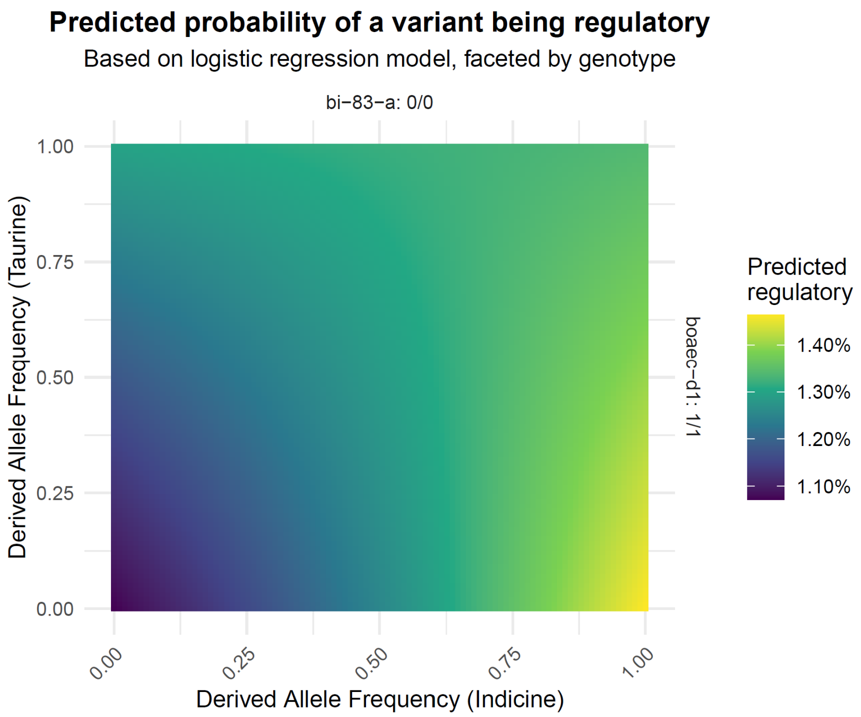


**Supplementary Figure 6.** Predicted probability of a variant being regulatory as a function of its Derived Allele Frequency (DAF) in Taurine and Indicine populations. Predictions are from the final logistic regression model (Supplementary Table 2). For the predictions in this plot the local variant density and PROCap levels were kept constant at their median levels and the Bi-83-A and BAOEC-D1 genotypes set to 0/0 and 1/1 respectively (homozygote reference and homozygote alternate). The plot illustrates the significant negative interaction between the two DAFs: the probability of being regulatory is highest (more yellow) when a variant's derived allele is common in one population but rare in the other, a pattern consistent with population-specific differentiation.

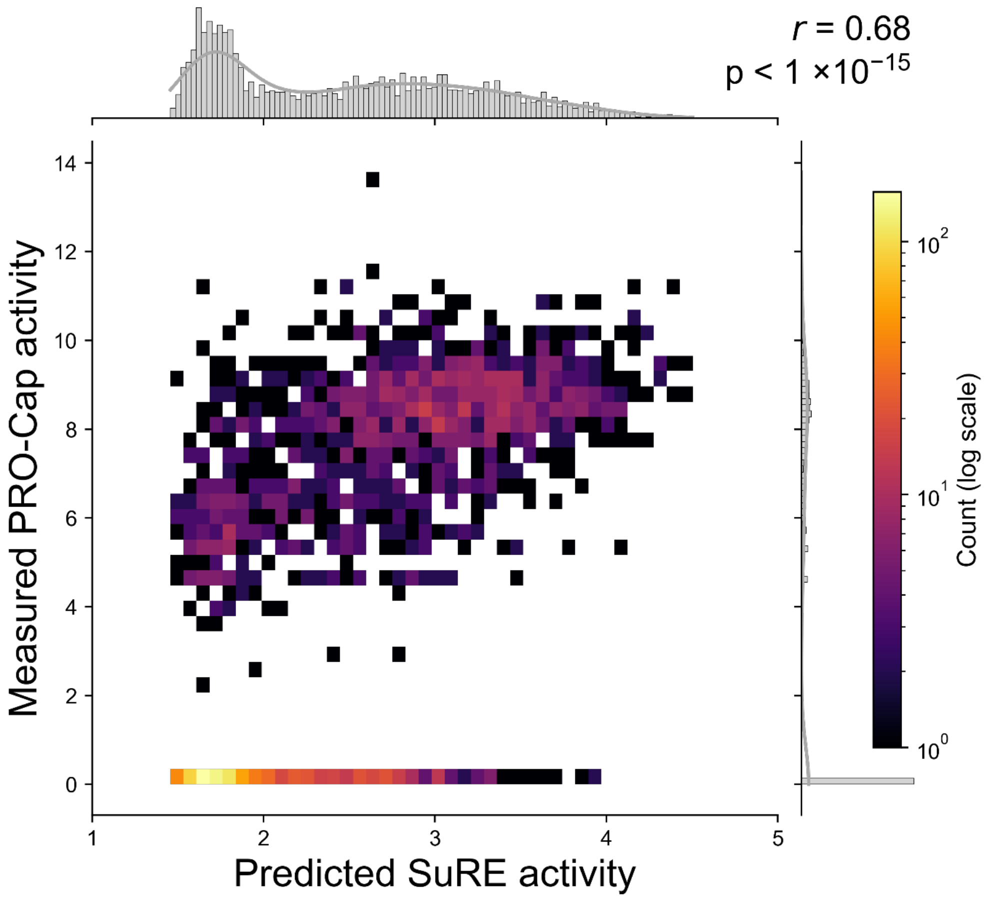


**Supplementary Figure 7**. EvaReg predictions against measure PRO-Cap scores. EvaReg model trained on cattle data has a positive and higher correlation with measured PRO-Cap data ($r=0.68$; $p<1\times10^{-15}$)

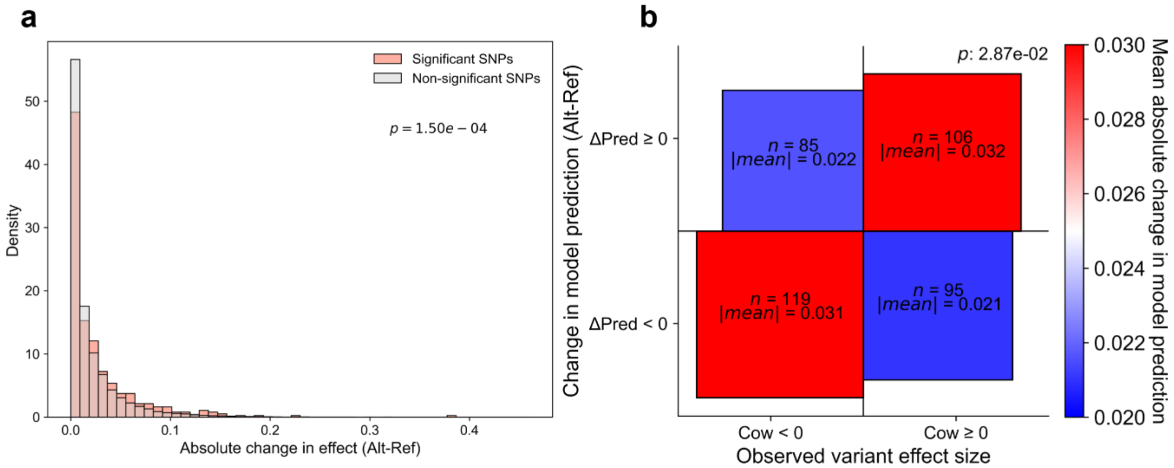


**Supplementary Figure 8**. EvaReg predicts variant effects in the SuRE dataset. (a) The absolute effect change in model predictions for a given SNP. A significant difference could be observed between the change in effects between significant SNPs and non-significant SNPs (Kolmogorov-Smirnov test; $p=1.5\times10^{-4}$). (b) Fourfold plot showing the change in direction of effect for a given SNP. A binomial test confirms a significant difference between correct and incorrect identification ($p=0.029$)

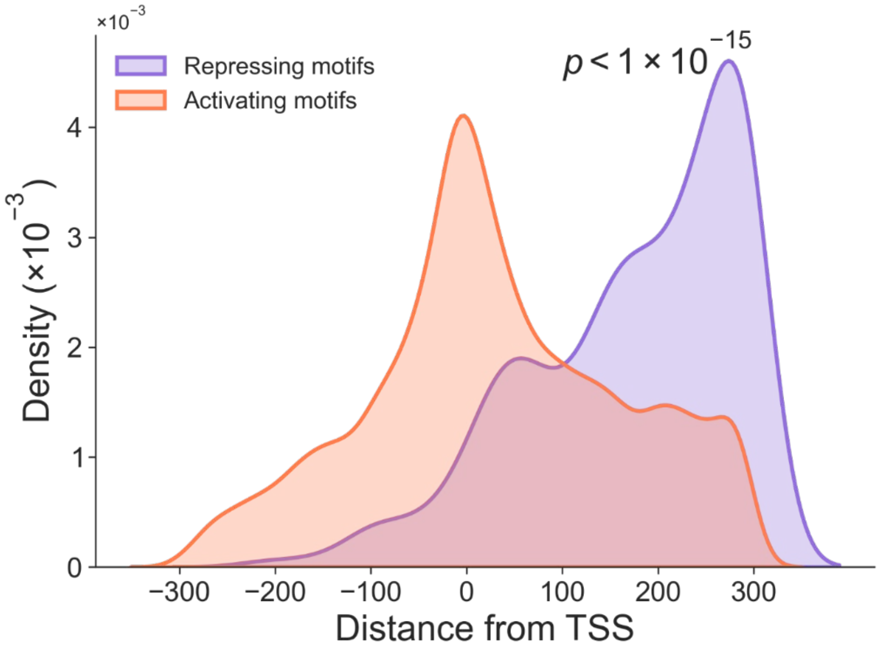


**Supplementary Figure 9**. Distribution of all motifs identified by EvaReg in the promoter dataset for the human SuRE data as activating (orange) or repressive (purple) (Kolmogorov–Smirnov test; $P<1\times10^{-15}$).

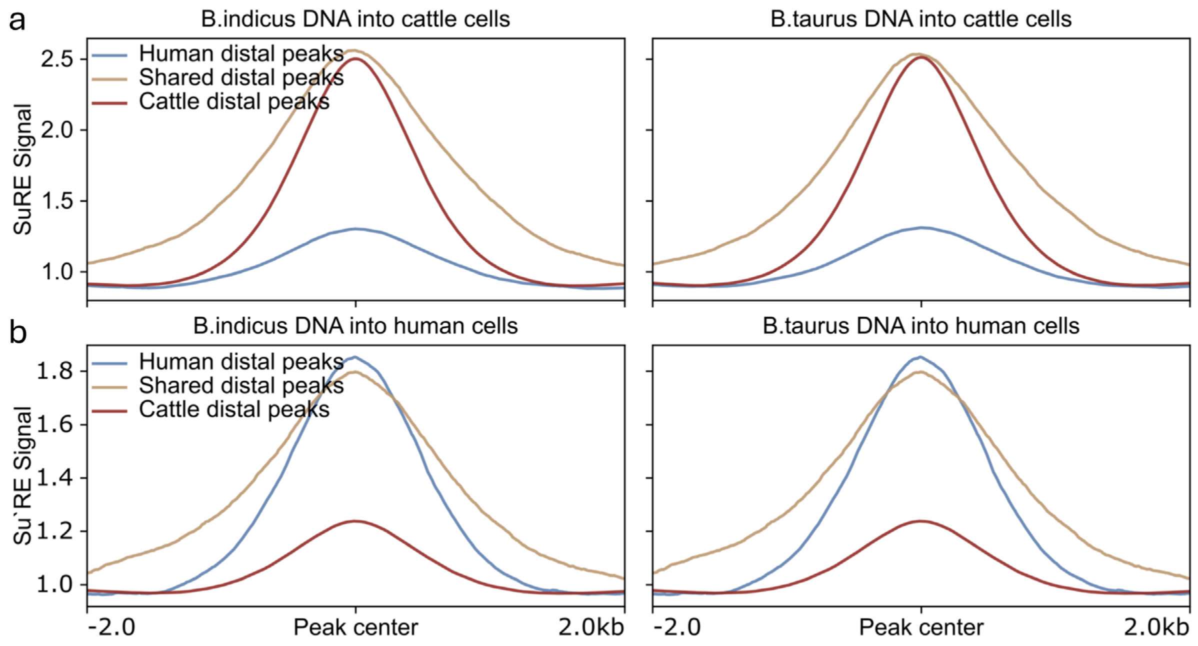

**Supplementary Figure 10.** Species-specific SuRE signal across distal regulatory peak sets. (a) Cattle SuRE signal centred on cattle-specific, human-specific, and shared distal peaks. Signals are shown for *B. indicus* (left) and *B. taurus* (right) DNA assayed in cattle cells. (b) Human SuRE signal centred on the same peak categories. Signals are shown for *B. indicus* (left) and *B. taurus* (right) DNA assayed in human cells.

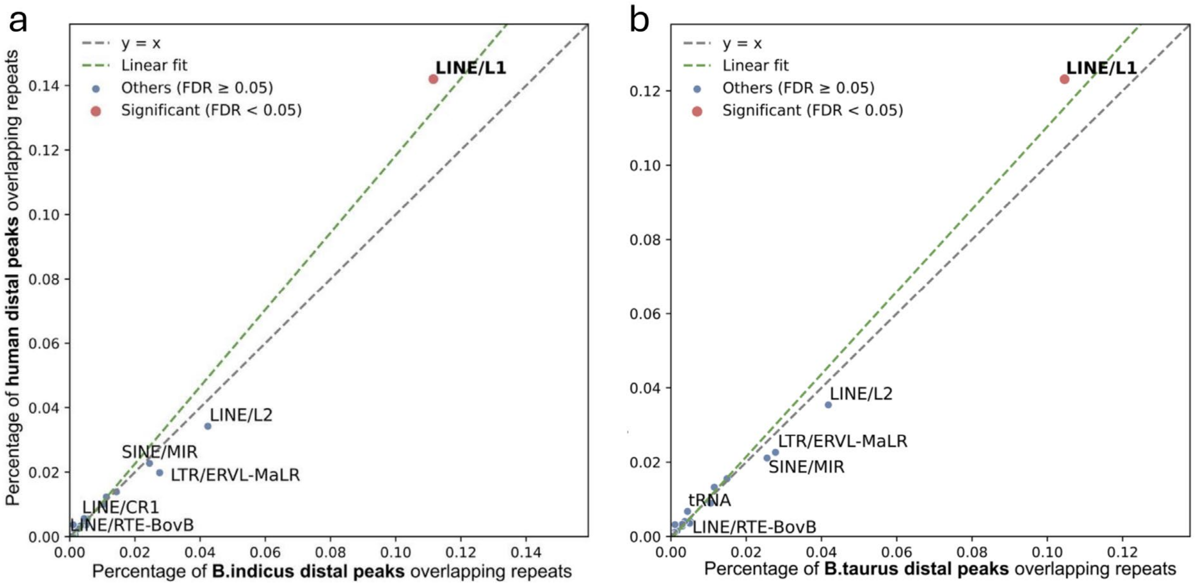


**Supplementary Figure 11.** Scatterplots show the percentage of distal peaks overlapping repeat classes in cattle and human datasets. The x-axis indicates the proportion of distal peaks from (a) *B. indicus* or (b) *B. taurus* overlapping each repeat class, and the y-axis indicates the corresponding proportion for human distal peaks. Each point represents a repeat class; labels are shown for classes that are significantly different between species (FDR < 0.05) or among those with the largest absolute differences in overlap between species. The dashed grey line denotes parity (x=y), and the dashed green line indicates the linear fit. Repeat classes with significantly different overlap between species are highlighted in red, and non-significant classes are shown in blue. LINE/L1 elements exhibit increased overlap in human cell-specific distal peaks compared to cattle cell-specific distal peaks.

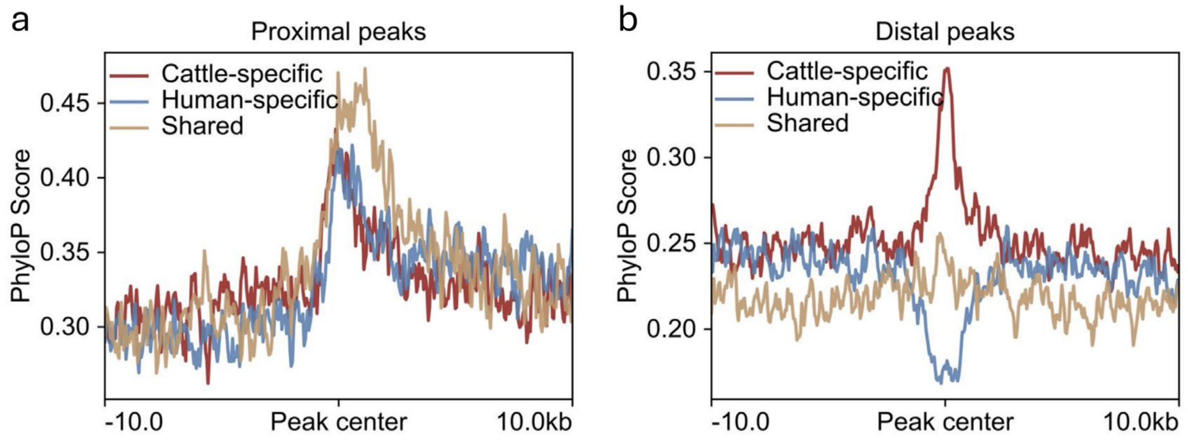


**Supplementary Figure 12.** Sequence conservation at (a) proximal and (b) distal peaks after removing repeat-overlapping regions. Signals are plotted relative to peak centres (±10 kb) for cattle-specific, human-specific, and shared peak sets.

### Supplementary Tables

| Tissue | Significant in Both | Significant in Cattle only | Significant in Human only |
|---|---|---|---|
| Adipose | 0.33 | 3.22E-29 | 0.96 |
| Blood | 0.58 | 1.41E-15 | 0.10 |
| Embryo | 0.31 | 3.36E-12 | 0.18 |
| Hypothalamus | 0.00 | 3.26E-15 | 0.69 |
| Intramuscular fat | 0.14 | 6.39E-24 | 0.98 |
| Jejunum | 0.37 | 6.17E-21 | 0.90 |
| Kidney | 0.83 | 8.16E-19 | 0.91 |
| Leukocyte | 0.34 | 1.27E-09 | 0.50 |
| Liver | 0.89 | 4.14E-09 | 0.84 |
| Lung | 0.10 | 2.38E-14 | 0.98 |
| Lymph node | 0.15 | 6.91E-25 | 0.95 |
| Macrophage | 0.94 | 3.07E-07 | 0.88 |
| Mammary | 0.34 | 1.52E-17 | 1.00 |
| Mammary L | 0.69 | 3.87E-13 | 0.37 |
| Milk cell | 0.08 | 3.81E-12 | 0.42 |
| Monocytes | 0.99 | 2.79E-10 | 0.23 |
| Muscle | 0.33 | 1.90E-15 | 1.00 |
| Muscle cross | 0.74 | 5.71E-23 | 0.71 |
| Muscle indicus | 0.22 | 2.69E-12 | 0.58 |
| Muscle taurus | 0.16 | 4.26E-09 | 1.00 |
| Ovary | 0.81 | 2.02E-11 | 0.31 |
| Oviduct | 0.02 | 1.36E-10 | 0.29 |
| Pituitary | 0.00 | 1.79E-06 | 0.62 |
| Rumen | 0.53 | 1.12E-11 | 0.77 |
| Spleen | 0.78 | 7.17E-04 | 0.01 |
| Testis | 0.43 | 5.37E-13 | 0.89 |
| Uterus | 0.53 | 1.34E-17 | 1.00 |

**Supplementary Table 1.** P values of enrichment of cattle GTEx eQTL signal among emVars, by tissue (Wilcoxon tests).

| Dependent variable | Odds ratio | 95% confidence interval | P value |
|---|---|---|---|
| DAF (Taurine) | 1.213 | (1.177, 1.249) | $<2\times10^{-16}$ |
| DAF (Indicine) | 1.371 | (1.332, 1.410) | $<2\times10^{-16}$ |
| DAF Taurine × DAF Indicine (interaction) | 0.753 | (0.684, 0.823) | $1.6\times10^{-15}$ |
| Local variant density (log-transformed) | 1.815 | (1.806, 1.823) | $<2\times10^{-16}$ |

| | | | |
|---|---|---|---|
| Local PROCap level (log-transformed) | 1.236 | (1.228, 1.245) | $<2x10^{-16}$ |
| B.Indicus sample genotype (0/0) | 0.836 | (0.805, 0.867) | $<2x10^{-16}$ |
| B.Indicus sample genotype (1/1) | 0.819 | (0.786, 0.853) | $<2x10^{-16}$ |
| B.Taurus sample genotype (0/0) | 0.895 | (0.868, 0.922) | $3.4x10^{-16}$ |
| B.Taurus sample genotype (1/1) | 0.915 | (0.872, 0.957) | $4.2x10^{-05}$ |
| B.Indicus genotype (1/1) × B.Taurus genotype (0/0) | 1.864 | (1.827, 1.900) | $<2x10^{-16}$ |
| B.Indicus genotype (0/0) × B.Taurus genotype (1/1) | 1.779 | (1.727, 1.831) | $<2x10^{-16}$ |
| Constant | 0.003 | (-0.032, 0.038) | $<2x10^{-16}$ |

**Supplementary Table 2.** Logistic regression model for predictors of regulatory activity**.** Odds ratios (ORs) and 95% confidence intervals from the binomial logistic regression model predicting whether a variant was linked to regulatory activity in cattle cells. Candidate predictors included derived allele frequency (DAF) in taurine and indicine cattle, local variant density score (the number of other variants within 500bp), the genotypes of the DNA source samples at the variant, local PROCap signal intensity (the highest PROCap read count within 500bp), and interaction terms. Coefficients are reported as exponentiated model estimates (odds ratios), such that OR > 1 indicates increased odds of regulatory activity and OR < 1 indicates decreased odds. Continuous predictors marked as log-transformed were fitted on the natural-log scale.

| | **Human** | | **Pig** | |
|---|---|---|---|---|
| **Peak category** | **Distal** | **Proximal** | **Distal** | **Proximal** |
| Cattle-specific | 52.19% | 60.84% | 52.19% | 62.09% |
| Human-specific | 33.37% | 49.18% | 32.98% | 49.86% |
| Shared | 44.72% | 60.02% | 45.28% | 60.84% |

**Supplementary Table 3.** The proportion of peaks in each category for which an orthologous region could be identified in the human or pig genomes.

| | ***Bos taurus* hap1** | ***Bos taurus* hap2** | ***Bos indicus* hap1** | ***Bos indicus* hap2** |
|---|---|---|---|---|
| # reads | 7024874 | | 5857845 | |
| Read N50 | 16961 | | 19073 | |
| Effective coverage | 38.99 | | 37.24 | |
| # contigs | 866 | 833 | 998 | 815 |
| Total contig length | 3,069,909,559 | 3,146,488,503 | 3,071,328,995 | 3,065,239,272 |

| | | | | |
|---|---|---|---|---|
| Average contig length | 3,544,930 | 3,777,297 | 3,077,484 | 3,761,030 |
| Contig N50 | 77,452,409 | 90,034,391 | 79,425,261 | 86,019,493 |
| Contig L50 | 15 | 14 | 15 | 14 |
| Largest contig | 135,901,988 | 159,613,024 | 156,649,277 | 156,646,979 |
| Smallest contig | 13,447 | 13,726 | 16,748 | 12,009 |
| GC content % | 43.69 | 44.12 | 43.6 | 43.84 |

**Supplementary Table 4.** Read and assembly metrics for genomes of source animals.

| Data | | Count |
|---|---|---|
| Bi-83-A library iPCR sequencing read pairs | | 2,111,100,122 |
| BAOEC-D1 library iPCR sequencing read pairs | | 2,043,979,288 |
| cDNA reads from fragments aligned to Bi-83-A in linear reference genome | Cow 1 | 150,470,359 |
| | Cow 2 | 164,631,209 |
| | Hum 1 | 110,058,144 |
| | Hum 2 | 114,668,824 |
| cDNA reads from fragments aligned to BAOEC-D1 in linear reference genome | Cow 1 | 158,723,197 |
| | Cow 2 | 173,730,380 |
| | Hum 1 | 116,263,203 |
| | Hum 2 | 135,332,235 |
| cDNA reads from fragments aligned to Bi-83-A in genome graph | Cow 1 | 150,644,870 |
| | Cow 2 | 164,456,153 |
| | Hum 1 | 112,793,693 |
| | Hum 2 | 118,032,627 |
| cDNA reads from fragments aligned to BAOEC-D1 in genome graph | Cow 1 | 153,922,171 |
| | Cow 2 | 168,310,639 |
| | Hum 1 | 116,410,788 |
| | Hum 2 | 136,386,515 |

**Supplementary Table 5**. Read and cDNA count statistics for the SuRE data.

| Sample ID | Accession | Subspecies |
|---|---|---|
| Brahman_BRAAUSM003 | SRR6650020 | *Bos indicus* |
| Brahman_BRAAUSM004 | SRR6650021 | *Bos indicus* |
| Brahman_BRAAUSM001 | SRR6650022 | *Bos indicus* |
| Brahman_BRAAUSM046 | SRR6650031 | *Bos indicus* |
| Gir_33534 | SRR4280088 | *Bos indicus* |
| Gir_33537 | SRR4280093 | *Bos indicus* |

| | | |
|---|---|---|
| Gir_33539 | SRR4280097 | *Bos indicus* |
| Gir_33540 | SRR4280102 | *Bos indicus* |
| Nelore_SAMN05788522 | SRR4280132 | *Bos indicus* |
| Nelore_SAMN05788524 | SRR4280142 | *Bos indicus* |
| Sahiwal_Sha3b | SRR6936540 | *Bos indicus* |
| Tharparkar_Thar1 | SRR6936538 | *Bos indicus* |
| Hariana_Har03 | SRR6936539 | *Bos indicus* |
| Ongole_1 | SRR19867309 | *Bos indicus* |
| Kasargod_Dwarf_2 | SRR19867308 | *Bos indicus* |
| Kasargod_Kapila_2 | SRR19867306 | *Bos indicus* |
| Vechur | SRR19867304 | *Bos indicus* |
| Gir_04 | ERR11600027 | *Bos indicus* |
| Gir_06 | ERR11605912 | *Bos indicus* |
| Gir_08 | ERR11604870 | *Bos indicus* |
| Gir_12 | ERR11605080 | *Bos indicus* |
| Gir_250 | ERR13488881 | *Bos indicus* |
| Gir_376 | ERR13502943 | *Bos indicus* |
| Gir_383 | ERR13448261 | *Bos indicus* |
| Gir_390 | ERR13459803 | *Bos indicus* |
| Gir_446 | ERR13492134 | *Bos indicus* |
| Gir_713 | ERR13461126 | *Bos indicus* |
| Hariana_11 | ERR13453568 | *Bos indicus* |
| Hariana_16 | ERR13488876 | *Bos indicus* |
| Hariana_26 | ERR13623139 | *Bos indicus* |
| Hariana_31 | ERR13448232 | *Bos indicus* |
| Hariana_39 | ERR13459804 | *Bos indicus* |
| Hariana_40 | ERR13459805 | *Bos indicus* |
| Kankrej_107 | ERR11635185 | *Bos indicus* |
| Kankrej_59 | ERR11606042 | *Bos indicus* |
| Ladakhi_199 | ERR11605898 | *Bos indicus* |
| Red_Sindhi_134 | ERR11635917 | *Bos indicus* |
| Red_Sindhi_135 | ERR11630203 | *Bos indicus* |
| Red_Sindhi_37 | ERR11606177 | *Bos indicus* |
| Red-Sindhi_38 | ERR11605602 | *Bos indicus* |
| Sahiwal_22 | ERR11605649 | *Bos indicus* |
| Sahiwal_24 | ERR11605854 | *Bos indicus* |
| Sahiwal_26 | ERR11602464 | *Bos indicus* |
| Sahiwal_27 | ERR11605987 | *Bos indicus* |
| Sahiwal_402 | ERR13502944 | *Bos indicus* |
| Sahiwal_605 | ERR13461119 | *Bos indicus* |
| Sahiwal_804 | ERR13461128 | *Bos indicus* |
| Sahiwal_825 | ERR13492154 | *Bos indicus* |
| Sahiwal_857 | ERR13488872 | *Bos indicus* |
| Tharparkar_04 | ERR13492137 | *Bos indicus* |
| Tharparkar_07 | ERR13461127 | *Bos indicus* |

| | | |
|---|---|---|
| Tharparkar_142 | ERR11607656 | *Bos indicus* |
| Tharparkar_143 | ERR11607666 | *Bos indicus* |
| Tharparkar_144 | ERR11607675 | *Bos indicus* |
| Tharparkar_145 | ERR11607309 | *Bos indicus* |
| Tharparkar_19 | ERR13488878 | *Bos indicus* |
| Angus_186 | SRR4279936 | *Bos taurus* |
| Angus_219 | SRR4279940 | *Bos taurus* |
| Angus_261 | SRR4279946 | *Bos taurus* |
| Angus_294 | SRR4279950 | *Bos taurus* |
| Angus_342 | SRR4279954 | *Bos taurus* |
| SwedishRed_SAMEA3871773 | ERR1256961 | *Bos taurus* |
| ScotHighland_HC26 | ERR1747025 | *Bos taurus* |
| ScotHighland_RM23 | ERR1747043 | *Bos taurus* |
| BrownSwiss_NDA002 | ERR1766310 | *Bos taurus* |
| BrownSwiss_DNA45641 | ERR1747023 | *Bos taurus* |
| BrownSwiss_RM721 | ERR1746314 | *Bos taurus* |
| BrownSwiss_RM723 | ERR1746315 | *Bos taurus* |
| BrownSwiss_RM763 | ERR1746326 | *Bos taurus* |
| Eastern-Finncattle_SAMEA4827182 | ERR2734942 | *Bos taurus* |
| Eastern-Finncattle_SAMEA4827183 | ERR2734943 | *Bos taurus* |
| Eastern-Finncattle_SAMEA4827184 | ERR2734944 | *Bos taurus* |
| Eastern-Finncattle_SAMEA4827185 | ERR2734945 | *Bos taurus* |
| Hereford_SAMN02843132 | SRR1365128 | *Bos taurus* |
| Hereford_SAMN02841110 | SRR1346384 | *Bos taurus* |
| Hereford_SAMN02843091 | SRR1365140 | *Bos taurus* |
| Hereford_SAMN02843109 | SRR1365142 | *Bos taurus* |
| Hereford_SAMN02843090 | SRR1365137 | *Bos taurus* |
| Charolais_SAMN02842704 | SRR1348574 | *Bos taurus* |
| Charolais_SAMN02842705 | SRR1348577 | *Bos taurus* |
| Charolais_SAMN02842706 | SRR1348578 | *Bos taurus* |
| Charolais_SAMN02843108 | SRR1365145 | *Bos taurus* |
| DabieshanCattle_AHWD1 | SRR5507192 | *Bos taurus* |
| Jersey_0067 | SRR7340696 | *Bos taurus* |
| Jersey_0062 | SRR7340716 | *Bos taurus* |
| Jersey_0029 | SRR7340745 | *Bos taurus* |
| Jersey_0023 | SRR7340772 | *Bos taurus* |
| Jersey_0024 | SRR7340779 | *Bos taurus* |
| Limousine_RM381 | ERR1766316 | *Bos taurus* |
| Limousine_7440 | SRR4280046 | *Bos taurus* |
| Limousine_SAMN02843095 | SRR1365134 | *Bos taurus* |
| Limousine_SAMN02843094 | SRR1365136 | *Bos taurus* |
| Romagnola_33844 | SRR4280155 | *Bos taurus* |

| | | |
|---|---|---|
| Simmenthal_71407 | SRR4280192 | *Bos taurus* |
| Simmenthal_71411 | SRR4280196 | *Bos taurus* |
| Simmenthal_71415 | SRR4280200 | *Bos taurus* |
| Simmenthal_71420 | SRR4280204 | *Bos taurus* |
| Simmenthal_71657 | SRR4280207 | *Bos taurus* |
| Western-Fincattle_SAMEA4827187 | ERR2734947 | *Bos taurus* |
| Western-Fincattle_SAMEA4827188 | ERR2734948 | *Bos taurus* |
| Western-Fincattle_SAMEA4827189 | ERR2734949 | *Bos taurus* |
| Western-Fincattle_SAMEA4827191 | ERR2734951 | *Bos taurus* |
| Holstein_1 | ERR2694921 | *Bos taurus* |
| Holstein_5 | ERR2694925 | *Bos taurus* |
| Holstein_23 | ERR2694943 | *Bos taurus* |
| Holstein_28 | ERR2694948 | *Bos taurus* |
| Holstein_32 | ERR2694952 | *Bos taurus* |
| Holstein_40 | ERR2694960 | *Bos taurus* |
| Holstein_45 | ERR2694965 | *Bos taurus* |
| Holstein_55 | ERR2694975 | *Bos taurus* |
| Holstein_64 | ERR2694984 | *Bos taurus* |
| Holstein_72 | ERR2694992 | *Bos taurus* |

**Supplementary Table 6**. The public samples used to calculate *B. indicus* and *B. taurus* allele frequencies.
